# Developmental Divergence of Sensory Stimulus Representation in Cortical Interneurons

**DOI:** 10.1101/2020.04.28.065680

**Authors:** Rahel Kastli, Rasmus Vighagen, Alexander van der Bourg, Ali Ozgur Argunsah, Asim Iqbal, Fabian F. Voigt, Daniel Kirschenbaum, Adriano Aguzzi, Fritjof Helmchen, Theofanis Karayannis

**Affiliations:** Laboratory of Neural Circuit Assembly, Brain Research Institute, University of Zurich Winterthurerstrasse 190, CH-8057 Zurich, Switzerland; Laboratory of Neural Circuit Dynamics, Brain Research Institute, University of Zurich Winterthurerstrasse 190, CH-8057 Zurich, Switzerland; Neuroscience Center Zurich, University of Zurich and ETH Zurich, Winterthurerstrasse 190, CH-8057 Zurich, Switzerland; Institute of Neuropathology, University Hospital Zurich, Schmelzbergstrasse 12, CH-8091 Zurich, Switzerland

## Abstract

Two inhibitory cell types involved in modulating barrel cortex activity and perception during active whisking in adult mice, are the VIP^+^ and SST^+^ interneurons. Here we identify a developmental transition point of structural and functional rearrangements onto these interneuron types around the start of active sensation at P14. Using in vivo two-photon Ca^2+^ imaging, we find that before P14, both interneuron types respond stronger to a multi-whisker stimulus, whereas after P14 their responses diverge, with VIP^+^ cells losing their multi-whisker preference and SST^+^ neurons enhancing theirs. Rabies virus tracings followed by tissue clearing, as well as photostimulation-coupled electrophysiology reveal that SST^+^ cells receive higher cross-barrel inputs compared to VIP^+^ at both time points. In addition, we also uncover that whereas prior to P14 both cell types receive direct input from the sensory thalamus, after P14 VIP^+^ cells show reduced inputs and SST^+^ cells largely shift to motor-related thalamic nuclei.

## Introduction

Postnatal development and maturation of neuronal circuits responsible for sensory processing is fundamental for accurate representation of the environment as animals transition into actively interacting with the external world^1^. Unravelling the mechanisms of neocortical development is key for understanding the emergence of purposeful and goal-directed motor actions informed by sensory cues. It is therefore of great importance to study the alterations that sensory cortical networks undergo in responding to diverse sensory stimuli upon developmental behavioural transitions.

The whisker primary somatosensory cortex (wS1) of rodents is a well-suited model to study sensory processing throughout development due to the somatotopic way in which the information is transmitted from the whiskers to the cortex, as well as its role in spatial navigation^2,3^. Although the whiskers of mice are already present at the time of birth, during the first postnatal weeks the whisker pad only exhibits spontaneous muscle twitches, which coincide with activity in wS1^4,5^. Around postnatal day 14 (P14), mice start displaying bilateral rhythmic movements of the whiskers (‘active whisking’), enabling them to explore their environment and extract more detailed information from their surrounding^6–8^.

Although the wS1 has been a subject of research for many decades, it is only recently that the role of inhibition in sensory processing in the adult cortex has started to be explored. Several studies showed that layer 2/3 (L2/3) vasoactive intestinal peptide-expressing (VIP^+^) interneurons (INs) in the wS1 can inhibit the inhibitory somatostatin-expressing (SST^+^) INs, thus leading to a net excitation of pyramidal cells (disinhibition)^9–11^. The same disinhibitory connectivity motif has also been described in other cortical areas, suggesting a general mechanism by which cortico-cortical loops can influence cortical processing^12–16^. In the wS1, this VIP^+^-SST^+^ disinhibitory loop can be recruited in a top-down fashion by the whisker primary motor cortex (wM1), which directly innervates the wS1 VIP^+^ INs in L2/3^9^. In addition it has been shown that VIP^+^ INs can be activated by bottom-up thalamic inputs, potentially also engaging in disinhibition through SST^+^ cells^17–19^. Nevertheless, a recent study reports opposite results, with SST^+^ cells being strongly activated by a whisker-driven sensory stimulus, compared to VIP^+^ cells which are found to be silenced^20^. We hypothesized that the main reason behind this discrepancy between the aforementioned results is the stimulus paradigm used. Researchers either stimulated one single whisker^11^ or multiple whiskers at the same time^20^-two paradigms that would engage the adult barrel cortex in very distinct ways. We further hypothesized that these interneurons would be very differently engaged by these stimulation paradigms prior to P14, when top-down modulation is absent and discrimination of fine features through the somatosensory system are probably not yet developed. When we tested these hypotheses, we indeed found that the strength of activation of VIP^+^ and SST^+^ INs depends on the nature of the presented stimulus. We find that compared to the single-whisker deflection, the multi-whisker stimulation leads to a higher activation of both cell types before P14, but after the onset of active whisking only SST^+^ exhibit higher activation. Intriguingly, these response alterations are accompanied by a significant rearrangement of thalamic connections onto both VIP^+^ and SST^+^ INs in the same time window, later than any other thalamo-cortical connectivity restructuring reported to date.

## Results

### Divergent sensory stimulus responses in superficial cortical VIP^+^ and SST^+^ interneurons during development

To assess how wS1 VIP^+^ and SST^+^ INs in L2/3 respond to whisker stimuli across development we performed acute *in vivo* two-photon calcium (Ca^2+^) imaging under light-anesthesia at two developmental time points; prior to the onset of active whisking (age: P8-12) and after the beginning of active whisking (age: P21-41, denoted as P21+). To visualize VIP^+^ and SST^+^ IN activity, we used animals expressing tdTomato in either VIP^+^ or SST^+^ INs (VIPCre-Ai14 and SSTCre-Ai14 lines, respectively) and injected the membrane-permeable AM-ester form of the Ca^2+^ indicator OGB-1 into wS1. Two-photon Ca^2+^ imaging was performed during spontaneous activity and upon stimulation of either the principal C2 whisker one time alone (single-whisker stimulation) or both the principal whisker and the majority of the macro vibrissae once together (multi-whisker stimulation) (Figure 1b) ^21^.

**Figure 1:**
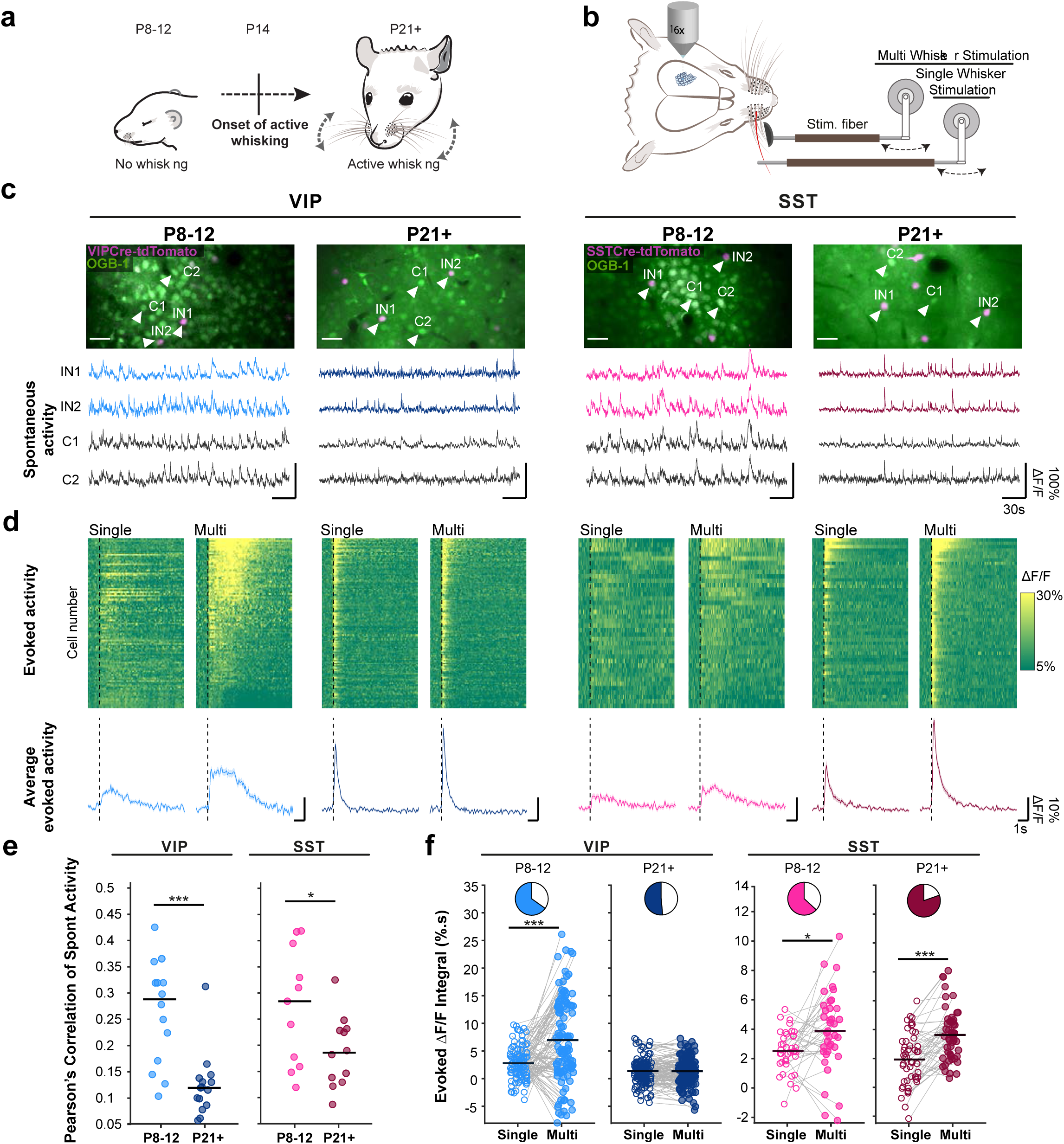
Divergent sensory stimulus responses in superficial cortical VIP+ and SST+ interneurons during development. **(a)** Acute in-vivo two-photon Ca^2+^ imaging was performed before (P8-12) and after (P21+) the start of active whisking. **(b)** Schematic represenation of whisker stimulation protocol. **(c)** Top: average intensity projection of calcium imaging regions after bulk loading of OGB-1 (green). lnterneurons are labeled with tdTomato using reporter mouse lines; scale bar 35 μm. Bottom: representative examples of raw ΔF/F traces of spontaneous activity of two interneurons (IN1,2) and two surrounding cells (C1,2). **(d)** Top: ΔF/F signal over time of all recorded VIP and SST cells after whisker stimulation. Ca^2+^ responses are baseline corrected and aligned to whisker stimulus onset (dashed line). Cells are sorted by their cumulative activity following multi-whisker stimulation. (N=3 animals per group, VIP P8-12:109 cells, SST P8-12: 38 cells, VIP P21+: 138 cells, SST P21+: 51 cells) Bottom: Average ΔF/F responses with SEM. **(e)** Pearson’s correlations of spontaneous neuronal activity within each interneuron type before and after P14. Same number of cells are included as in d) but the data is plotted by imaging spot. **(f)** Average of the evoked ΔF/F integral after single and multi whisker stimulation. Pie charts show the fraction of cells that increase (coloured) or decrease (white) in activity by multi vs. single whisker stimulation (N=3 animals per group, VIP P8-12:109 cells, SST P8-12: 38 cells, VIP P21+: 138 cells SST P21+: 51 cells). Statistics: e) Mann–Whitney U test, f) paired Wilcoxon-signed-rank test (*p<0.05, **p<0.01, ***p<0.001).

We first tested whether VIP^+^ and SST^+^ INs are functionally integrated before P14 by examining their spontaneous activity. We observed prominent Ca^2+^ transients in both IN types that correlated with the activity of the surrounding cells (Figure 1c, **Figure S1a**), indicating that they receive input capable of driving action potentials before the onset of active whisking. Previous studies have shown that cortical pyramidal neurons and superficial 5HT3a^+^ INs display highly correlated spontaneous activity within their population in early postnatal development and then undergo a de-correlation upon maturation of the circuit ^21–24^. To test if the same is true for VIP^+^ and SST^+^ INs we compared the correlations of spontaneous activity within each IN population before and after P14 and found a significant drop of co-activation with time (Figure 1e). Interestingly, at P21+, spontaneous activity of SST^+^ cells is more correlated than that of VIP^+^ cells, a phenomenon that has also been described in the visual cortex of adult mice (p<0.01, comparison not shown in the figure) ^25^. In addition, we compared the average correlation between SST^+^ and VIP^+^ INs and between the IN subtypes and their surrounding non-labeled cells, most likely excitatory neurons, again finding a similar decorrelation (**Figure S1a**).

Having established that VIP^+^ and SST^+^ INs are embedded in the developing circuit prior to P14, we next tested if they respond to whisker stimulation and if yes, how this would potentially change across development (Figure 1b). Since in a natural environment, both single and multiple whiskers can be deflected, we used both a single- and a multi whisker stimulus in both age groups, with the latter being even more relevant for neonatal mice, whose whiskers are physically closer together^5^.

As the response to a single-whisker stimulus in adult L2/3 VIP^+^ and SST^+^ INs has been described before by cell-attached electrophysiological recordings^17^, we first compared the activity of the two cell types in response to this stimulation paradigm. We found that at P21+ the peak ΔF/F of the average Ca^2+^ response in SST^+^ INs was lower compared to that of VIP^+^ INs, matching the published data^17^ (**Figure S1b**). Nevertheless, when looking at the integral of ΔF/F average response (over 8s following whisker stimulation) there was no significant difference between the two cell types (**Figure S1b**), and the same was true before P14 (**Figure S1c**).

We then assessed how IN responses to multi-compared to single-whisker stimulation may change during development by comparing the ΔF/F integral as an overall measure of direct or indirect activation of the cells by the sensory stimuli. The analysis showed that before the onset of active whisking (P8-12), both VIP^+^ and SST^+^ INs responded significantly stronger to multi-compared to single-whisker stimulation (Figure 1d, f). For the SST^+^ INs this continued to be the case and even enhanced after the onset of active whisking (P21+), while for the VIP^+^ INs the ΔF/F integral became similar between multi-versus single-whisker stimulation (Figure 1f). Since the Ca^2+^ transients recorded at P8-12 had a clear late component and a lower peak amplitude than the mature cells, we sought out to better understand how they relate to action potentials across development. We therefore filled SST^+^ INs with OGB-1 *in vitro* and evoked a set number of action potentials, while recording the Ca^2+^ transients in the cell bodies using two-photon microscopy. We found, that when comparing the ΔF/F responses between the two age groups, there was no significant difference detected, indicating that the integral of the Ca^2+^ responses can provide a good estimate for the functional activation of the cell types across development. Our results also suggest that the pronounced late component of the Ca^2+^ signal at P8-12 is not due to the differential Ca^2+^ buffering capacities at the two age points, but rather due to the way the network activates the cells (**Figure S1d**).

To address if the sensory-evoked Ca^2+^ responses we observed for the cell types and stimuli across development carry information that can make them discriminatory, we trained three different decoders to quantify differences in response profiles. Each decoder was trained on a subset of Ca^2+^ responses, evoked through either multi- or single-whisker stimulation, for each IN type and age. All of the classifiers showed above chance level values with VIPs displaying a reduction with age, whereas the SSTs an increase. This finding lends further support for the developmental divergence of these two cell type’s activation and function (**Figure S1e**).

### Superficial VIP^+^ and SST^+^ interneurons show distinct barrel-field afferent connectivity motifs

Increased responsiveness to multi-over single-whisker stimulation at both ages could be due to cortico-cortical inputs originating from surrounding barrel columns, which would provide convergent excitation originating from multiple whiskers. We therefore aimed at investigating the presynaptic inputs onto L2/3 VIP^+^ and SST^+^ INs at the two developmental time points, using a monosynaptic rabies virus approach^26^. This allowed us to assess both local and long-range connectivity of the two IN types.

By utilizing compound mouse genetics (VIPCre-HTB or SSTCre-HTB), in combination with the pseudo-typed rabies virus, direct presynaptic partners of L2/3 VIP^+^ and SST^+^ INs were labeled with mCherry. The primary infected cells (starter cells) could be identified by double labeling of mCherry and eGFP (provided by the HTB line). The viral injections were done at either P5 or P15 and after 7 days the brains were collected (at P12 and P22 respectively). A CLARITY-based tissue clearing protocol was performed on the brains and they were subsequently imaged in their entirety using a custom-built light-sheet microscope (mesoSPIM) (Figure 2a)^27^. The auto-fluorescence of the barrels allowed for detection of the barrel field and enabled us to accurately localize our injection site within the wS1 (Figure 2b-e and **Figure S2**). A second batch of injected brains was processed using classical histology, followed by eGFP immunostaining and widefield imaging. Both the cleared whole brain and the histology dataset were used to quantify the number and laminar position of the starter and the presynaptic cells located around the wS1 injection site. This dual approach was performed to safeguard that no information was lost while clearing the brains. For the histology dataset, a subset of all cut sections and a manual approach was used to identify the cells, whereas in the cleared brains a deep-neural network was trained to detect and count cells^28^. For both methods, the results were highly comparable (**Figure S3**).

**Figure 2:**
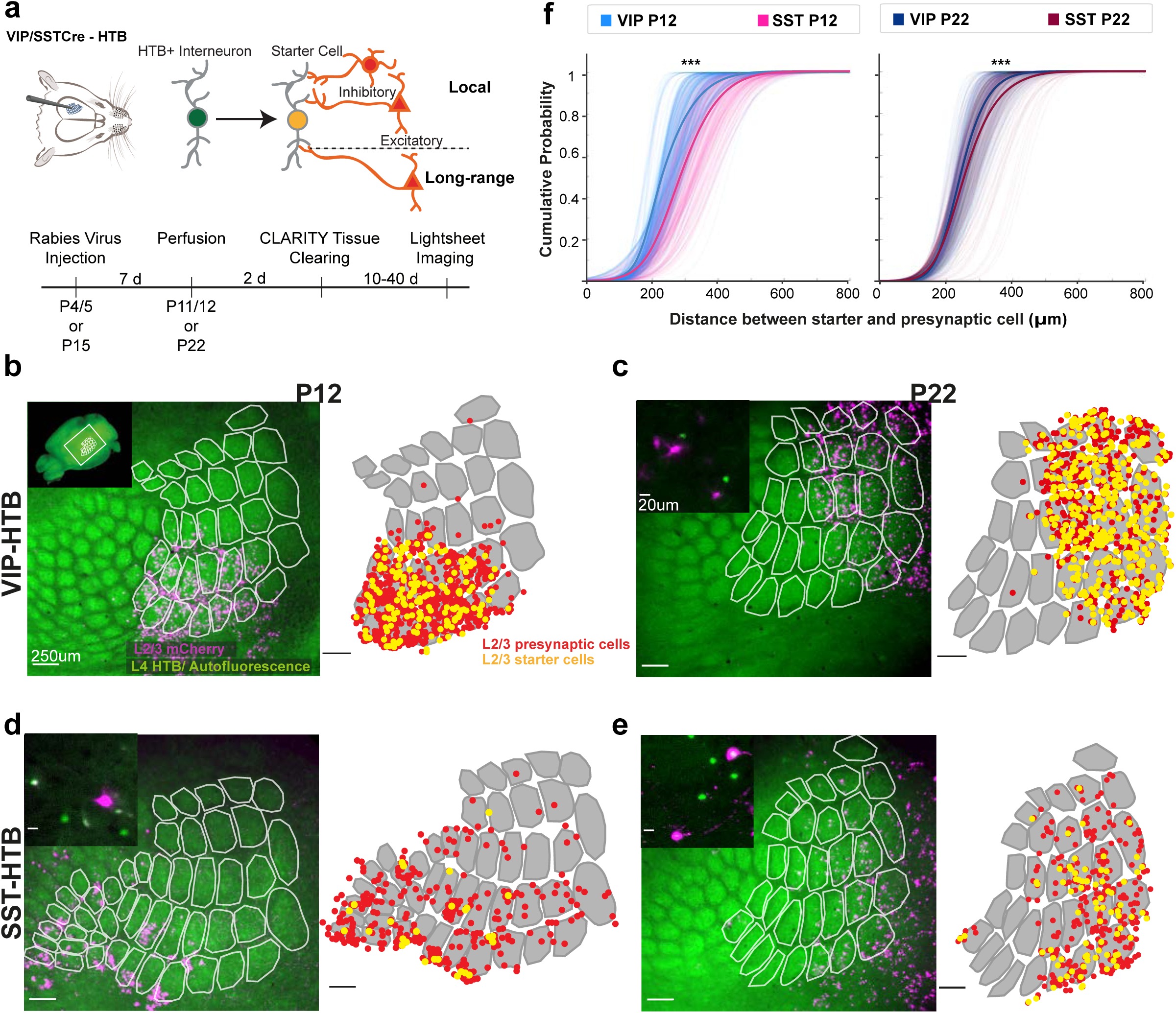
Superficial VIP+ and SST+ interneurons show distinct barrel-field afferent connectivity motifs. **(a)** Shematic representation of the experimental protocol. Rabies virus tracing was combined with tissue clearing and whole brain imaging **(b-e)** Left: Overlay of maximum intensity projection of L2/3 mCherry signal and median intensity projection of L4 autofluorescence after re-slicing of whole brain images. Right: Tranformation applied before distance analysis. Segmented barrels overlayed with L2/3 starter (yellow) and presynaptic (red) cells represented as dots. Inset in (b) indicates location of barrel field in the whole brain. Insets in (c-e) show close-ups of rabies labelled neurons. **(f)** Cumulative distribution of Euclidian distance in 2D between the starter cells and all presynaptic cells in a 800 μm radius around them. The faint lines in the background depict the distance distribution of each starter cell; the thicker lines depict the average. (VIP P12, N=4: 382 starter & 1943 presynaptic cells, SST P12, N=6: 102 starter & 581 presynaptic cells, VIP P22, N=5: 645 starter & 1561 presynaptic cells, SST P22, N=4: 110 starter & 504 presynaptic cells). Statistics: Kolmogorov-Smirnov Test (***p<0.001).

The analysis showed that although we see starter cells in L2-6 at both time points we get robust primary infection in L2/3 for both VIP^+^ and SST^+^ cells (**Figure S3a, b**). The distribution of the presynaptic cells across the layers mirrored that of the starter cells, with the VIP^+^ INs receiving more inputs from the upper cortical layers while the SST^+^ neurons more from the deeper layers (**Figure S3c, d**).

To assess if L2/3 SST^+^ and VIP^+^ INs receive a significant proportion of their inputs from neighboring barrels, the cleared 3D brain images were used to localize both starter and presynaptic cells within the barrel field (Figure 2b-e, **Figure S2a, b**). Only L2/3 presynaptic neurons located within wS1 were included in the analysis, since it is most likely that these inter-columnar inputs would originate from the upper layers^29,30^. As it is not possible to determine which presynaptic partner connects to which starter cell, a probabilistic approach was taken, where the distance between every starter to every presynaptic cell within 800µm distance was calculated and compared between the two cell types (**Figure S4**). The analysis revealed that SST^+^, compared to VIP^+^ INs, receive on average more distant inputs, both before and after P14 (Figure 2f). To further validate the results and overcome the variability observed in the data we also calculated the cumulative distance distribution of randomly selected starter and presynaptic cells (**Figure S2c)** (Online Methods). This anatomical data can help explain why the SST^+^ cells are able to report multi-over single-whisker activation both before and after P14. However, it cannot explain why the VIP^+^ INs are able to differentiate between multi- and single-whisker stimulation before P14 and not after, unless there is a change in the functional connectivity.

### Distinct functional connectivity motifs onto superficial VIP^+^ and SST^+^ interneurons

After revealing the anatomical connections of the wS1 cortical inputs onto superficial VIP^+^ and SST^+^ INs, we assessed their functionality and strength. Specifically, we investigated if after P14, SST^+^ INs receive stronger functional input from lateral sources compared to VIP^+^ INs, while before P14 they display more functionally similar input.

In order to investigate the developmental trajectory of VIP^+^ and SST^+^ IN intrinsic properties we performed whole-cell current-clamp recordings of L2/3 VIP^+^ or SST^+^ INs and analyzed their passive and active electrophysiological membrane properties (Figure 3b, **Figure S5a-e**). As expected, we found evidence of cell maturation in both cell types, with the membrane resistance decreasing and the action potentials becoming faster (**Figure S5a, d, e**). Nevertheless, the threshold for action potential generation dropped and action potential amplitude increased only in VIP^+^ INs, (**Figure S5b, c**), suggesting that the SST^+^ cells are closer to their mature state at P8-12.

**Figure 3:**
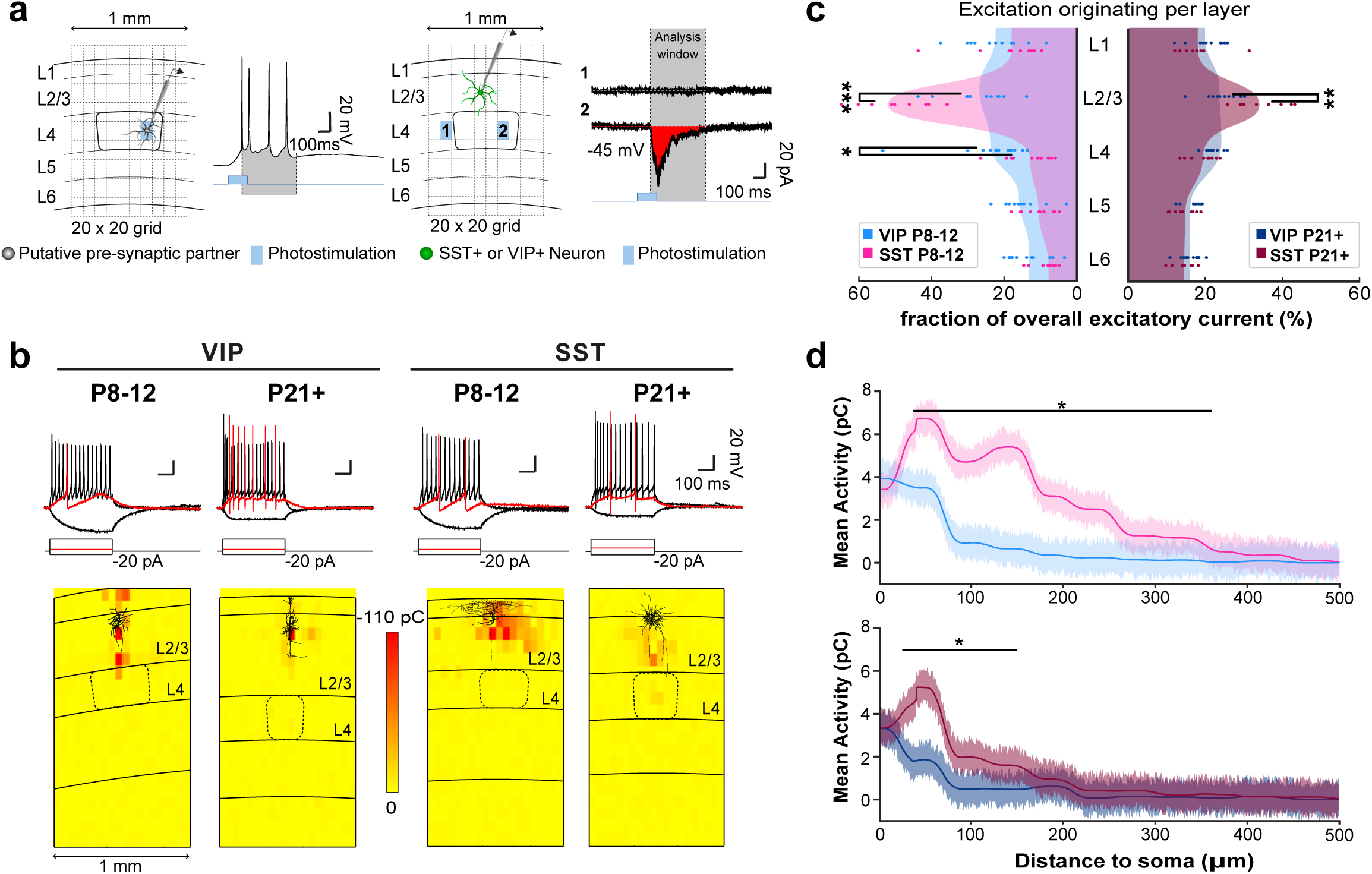
Distinct functional connectivity motifs onto superficial VIP+ and SST+ cortical interneurons. **(a)** Left: A graphic representation of photostimulation calibration with a patched putative pre-synaptic neuron within a neocortical slice indicated with layer boundaries (L1-6). Illumination calibration were done so that the paradigm successfully evoked a short train of APs in the putative pre-synaptic partners. AP discharge is displayed to the right, with the peak of the first AP until after the last AP creating a time-window used for future analysis of evoked currents. Right: schematic representation of photostimulation-based mapping of incoming currents onto interneurons through whole-cell patch-clamp recordings. Overlaid is a schematic of the 20×20 grid, indicating the quadrants that can be photostimulated. Two of the AP discharges of the interneuron is shown to the right, with one not eliciting (1) and one eliciting (2) a post-synaptic response. The time-window acquired during calibration is used to register and analyse the incoming currents. **(b)** Top: example traces of voltage responses to square hyperpolarizing and depolarizing current pulses, Δ +20pA. Bottom: Heatplot representations of normalized evoked excitatory current integral (in pC), recorded over development (P8-12 and p21+) from VIP+ (left) and SST+ (right) interneurons while performing glutamate uncaging in a grid pattern. **(c)** Plot of excitatory input onto VIP+ and SST+ cells (individual dots) averaged per lamina and normalized to average overall excitation within the field of view. The grand average of all cells per group is depicted as a continuous filled wave and compared within age groups. One data point has been excluded for illustration purposes only. **(d)** Mean evoked excitation originating from within L1-3 and plotted as a function of lateral distance from either side of the recorded cells somata. (VIP P8-12: 10 cells, SST P8-12: 10 cells, VIP P21+: 10 cells SST P21+: 9 cells) Statistics: Mann-Whitney U test for c) and two tailed t-test for d) (*p<0.05, **p<0.01, ***p<0.001)

In order to analyze the distribution of neurons providing synaptic input to the two IN types, we performed glutamate uncaging. This was carried out in a grid pattern onto wS1-containing brain slices while measuring evoked postsynaptic excitatory and inhibitory currents in L2/3 VIP^+^ or SST^+^ INs through whole-cell voltage-clamp recordings at −45mV (Figure 3a). With this approach, we mapped out the origin as well as the strength of incoming cortical connections at the two developmental stages investigated (P8-12 and P21+). We found that L2/3 SST^+^ INs receive the majority of their excitatory input from within their own layer and that this is the case both before and after P14. These results fit well with the laminar patterns of excitatory inputs described for the L2/3 SST/CR^+^ cells in adult mice^30^. The VIP^+^ cells on the other hand receive almost equal amounts of excitatory inputs from all layers, with a decrease in L5/6 (Figure 3c). Because the photostimulation was carried out over a large field of view we were able to assess lateral activity originating from up to 500 µm away from either side of the IN cell body and therefore inputs coming from outside the resident column. To compare our glutamate uncaging results to the rabies virus mapping, inputs located in the superficial layers (L1-3) were plotted in relation to their lateral distance from the recorded cell. In agreement with the rabies virus findings, the analysis showed that prior to P14 SST^+^ INs receive excitatory inputs from more lateral sources compared to VIP^+^ cells (Figure 3d). Even though this difference is reduced after P14, SST^+^ INs still receive a higher proportion of excitatory inputs from further away compared to VIP^+^ INs (Figure 3d). Interestingly, the VIP^+^ INs receive more inhibition than the SST^+^ INs from L2/3 both before and after P14, acting almost in an opposite fashion to the mapped incoming excitation (**Figure S5f, g**). Because glutamate uncaging could activate the receptors on the patched cell directly, and holding the cell at −45mV does not allow for a proper estimation of purely excitatory responses, a new set of photostimulation experiments was carried out in voltage-clamp mode at −70 mV, first without and then in the presence of tetrodotoxin (TTX). By subtracting the direct responses (in the presence of TTX) from the evoked excitatory ones, recorded before the addition of the drug, currents evoked exclusively from pre-synaptic neurons could be calculated (**Figure S5h, i**). The analysis of this new dataset showed that the SST^+^ INs do indeed receive excitation from more distant sources compared to the VIP^+^ cells within the superficial layers (Figure 3d).

The anatomical tracing data together with the functional incoming excitation data suggest that SST^+^ INs receive more lateral excitation compared to VIP^+^ cells at both developmental time points. This provides a plausible mechanism for the stronger activation of SST^+^ INs by multi-whisker stimulation. In contrast, despite that VIP^+^ cells also respond stronger to multi-compared to single-whisker stimuli before P14, no local cortical motif was detected that could help explain it. This suggests that the multi-whisker preference of VIP^+^ cells could be enabled by activity originating from long-range thalamic sources.

### Rearrangement of thalamic inputs onto superficial VIP^+^ and SST^+^ interneurons during development

Since the observed inter-barrel connectivity motifs are not able to fully explain the developmental changes in single-versus multi-whisker responses seen in the *in vivo* Ca^2+^ imaging data, we turned our attention to bottom-up inputs coming from the thalamus, labeled by our rabies tracings. The analysis was facilitated by the preservation of the 3D architecture of the thalamus in the cleared whole brain dataset. Interestingly, we found that before P14, both VIP^+^ and SST^+^ INs had many presynaptic cells in the thalamus (Figure 4a,c, **Movie S1, Movie S3**). The autofluorescence of the samples allowed us to distinguish between different thalamic nuclei; especially the ventral posteromedial nucleus of the thalamus (VPM), the primary relay station to the wS1, which was easily identifiable due to its distinct barreloid structure. Using this nucleus as a reference, we found that the thalamic areas providing inputs to superficial VIP^+^ and SST^+^ cells before P14 are mainly the VPM and the posterior complex (PO), a higher order nucleus primarily innervating wS1 L1-3 and L5a, as well as wS2^31^ (Figure 4a, c, e, **Movie S5**, **Movie S7**). Unexpectedly, we found a clear shift in the thalamic nuclei providing input to the SST^+^ INs after P14. In contrast to prior P14, very few or no pre-synaptic partners were found in the VPM or PO, while many cells were labeled in the Ventro-Medial nucleus (VM) and the Ventro-Anterior nucleus (VAL) of the thalamus (Figure 4d, e, **Figure S7**, **Movie S4**, **Movie S8**). Interestingly, after P14 the VIP^+^ INs did not seem to have any pre-synaptic partners in the thalamus (Figure 4b, **Movie S2, Movie S6**). This is in contrast to studies that have shown functional input from the thalamus onto VIP^+^ cells in the adult cortex^18,19,32^. Since it is widely known that the rabies virus does not spread to all the pre-synaptic partners of a cell^33–35^, and in our case this could also be augmented by reduced expression of the glycoprotein from the HTB mouse line^36^, we believe that this result reflects a reduction in thalamic inputs rather than a complete loss. To examine this, we used two alternative approaches. First, as an alternative to the HTB mouse line, an AAV HTB helper virus was co-injected with the rabies virus at P15 in VIPCre mice. The animals were sacrificed at P22 and the brains cleared and imaged as described above (n=3). Using this approach, we indeed detected a low number of pre-synaptic cells in the VPM and the PO (**Figure S7**). The second approach was to stain wS1-containing sections from VIPCre-tdTomato and SSTCre-tdTomato mice with the vesicular glutamate transporter 2 (VGlut2), a marker for thalamo-cortical terminals. This approach provides an independent validation of direct thalamo-cortical innervation onto superficially located VIP^+^ and SST^+^ INs, and in addition provides a measure of the strength of those inputs by means of number of synapses. To quantify the number of appositions, a custom-written program was used to analyze high-resolution confocal images (Online Methods). The analysis showed that prior to P14, VIP^+^ INs had significantly more VGlut2 puncta than SST^+^ INs, whereas after P14 the number of appositions onto VIP^+^ INs dropped significantly (Figure 4f), evening out the difference between the two cell types. These histological results support the data obtained with the rabies-based mapping, and suggest that upon the onset of active whisking thalamic input onto VIP^+^ INs is strongly reduced.

**Figure 4:**
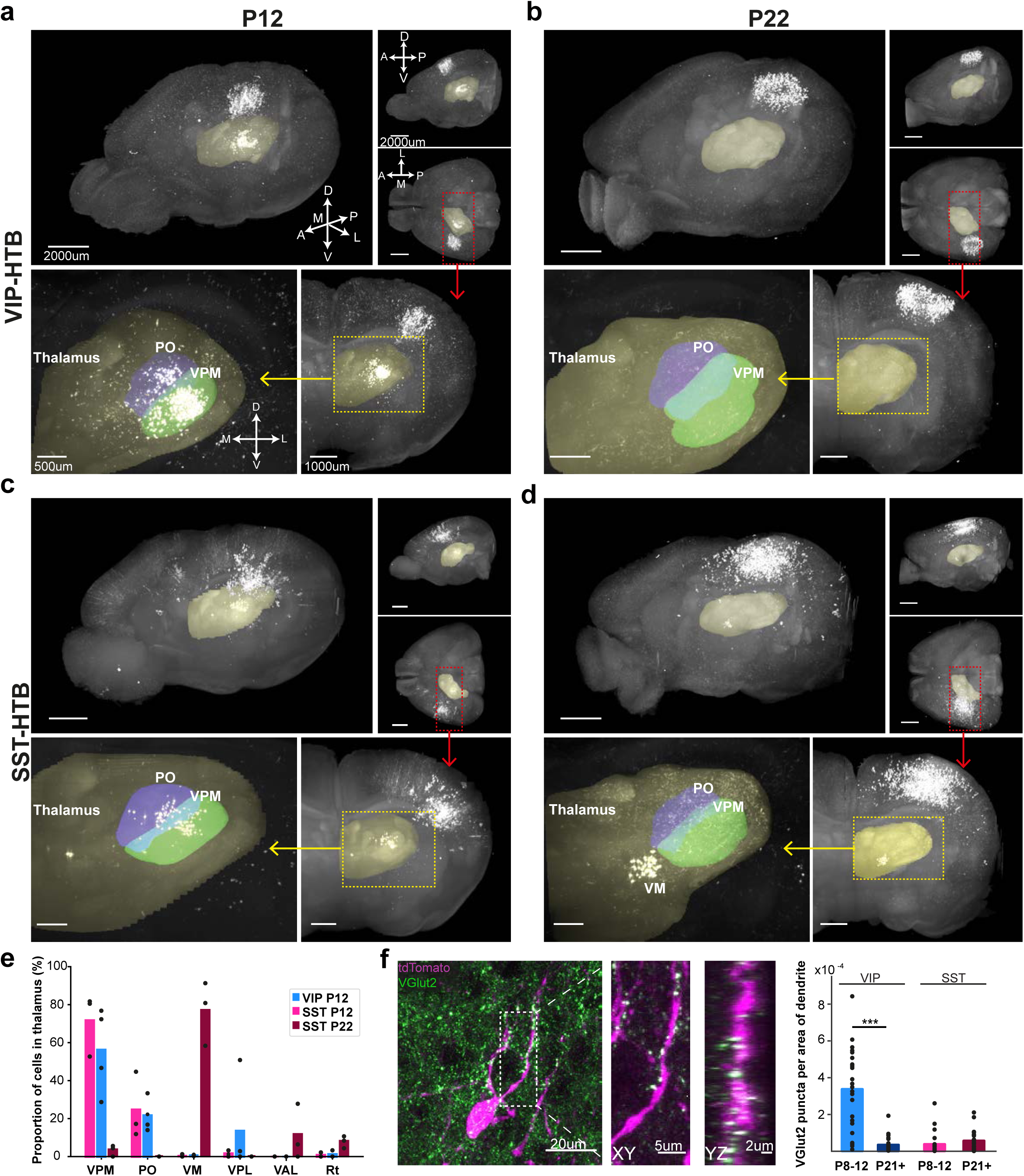
Rearrangement of thalamic inputs onto superficial VIP+ and SST+ interneurons during development. **(a-d)** Top panel: representative examples of cleared rabies injected brains seen from a oblique, top-down and side angle. The thalamus is highlighte in yellow. Red square indicates the area projected in bottom panel. Bottom panel: Coronal view of maximum intensity projection of the thalamus. Left: Overview. Right: zoom-in with VPM (green) and PO (blue) highlighted. **(e)** Quantification of pre-synaptic cells in different thalamic nuclei normalized to the total number of cells in the thalamus. VIP P22 is not included because no cells were found in the thalamus (N=3 brains for SST; N=4 brains for VIP P12). **(f)** Example of VGlut2 staining (green) in L2/3 of a wS1 section from a VIPCre-tdtomato animal at P9. The dotted square indicates the zoomed in part on the right, which shows a close-up of puncta appositions in xy and yz direction. Graph shows quantification of VGlut2 puncta on tdtomato-positive dendrites. The number of appositions in every picture is normalized to the area of tdtomato positive dendrites in the same picture. (N=3 brains per group; SST P21+: 28 images, rest: 26 images. Statistics: Mann–Whitney U test (***p<0.001).

Overall, our data suggest a model by which the VIP^+^ IN preference for multi-whisker stimulation before P14 is mainly supported by strong thalamo-cortical inputs that come from both VPM and PO. These inputs are significantly reduced after the onset of active whisking, together with the VIP^+^ INs response to multi-versus single-whisker stimulation (Figure 5). For the SST^+^ INs on the other hand, the preference for multi-whisker stimulation, both before and after P14 would be supported by the strong lateral inter-barrel connectivity, with a contribution of direct VPM and PO-derived excitation prior to P14 (Figure 5).

**Figure 5:**
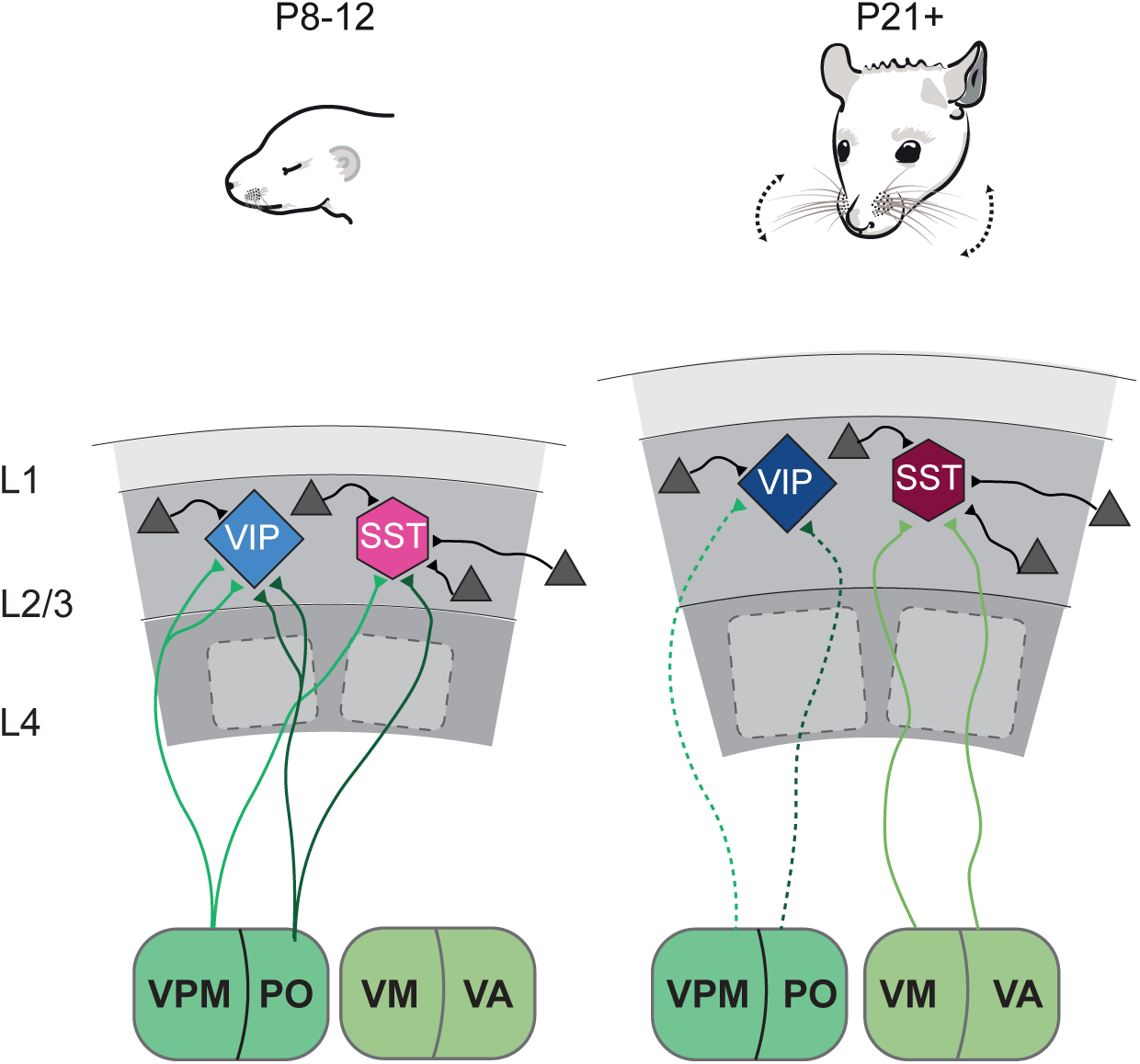
Schematic of developmental input rearrangement onto superficial VIP+ and SST+ interneurons. Local connectivity: SST+ interneurons receive exciatory inputs (grey triangles) from more distal source within the barrel cortex than VIP+ cells both before and after P14. Bottom-up inputs: Before P14 both VIP+ and SST+ cells receive the majority of their thalamic inputs from the VPM and PO nuclei of the thalamus, with the VIP+ cells showing more thalamic synapses on their dendrites. After P14 thalamic input onto VIP+ cells is strongly reduced while the input onto SST+ interneurons is shifted to the VM and VA thalamic nuclei.

## Discussion

This study investigates the engagement of VIP^+^ and SST^+^ INs by sensory stimuli, and the underlying circuits they are embedded in before and after the onset of active whisking (P14).

We initially hypothesized that in the juvenile mouse wS1, VIP^+^ and SST^+^ cells would be differentially activated by a single-versus a multi-whisker deflection respectively, based also on published reports^11,20^. By testing the activation pattern of the same population of INs for the two types of stimuli, we found support for our hypothesis. Juvenile VIP^+^ INs responded in the same manner whether a single or multiple whiskers were deflected, even though prior to P14 they responded much stronger to the multi-whisker stimulus. Juvenile SST^+^ cells on the other hand responded stronger to the multi-whisker stimulus, which was also the case before P14. Even though in this study we have not directly examined the connectivity between these two interneuron types, recently published work in the adult mice showed strong layer-dependent effects after VIP^+^ neuron Channelrhodopsin-based activation^20^. Specifically, the authors reported that L2-4 excitatory neurons did not increase their firing rate upon VIP^+^ neuron activation, suggesting that the recruitment of the VIP-SST inhibitory loop is layer- and most likely also stimulus-dependent. On the other hand the same study suggests that upon active whisking and stimulation of multiple whiskers, SST^+^ cells are strongly activated, whereas VIP^+^ INs are inhibited, with a delay that matches the activation of SST^+^ neurons. These results are in line with the reciprocity of the connections between the two IN types^12^, and together with our data suggest that the di-synaptic disinhibitory VIP-SST connection is engaged in a dynamic manner, with one cell inhibiting the other, the directionality of which is context-dependent.

Developmentally, the robust activation of both VIP^+^ and SST^+^ INs prior to P14 by the multi-compared to the single-whisker stimulus, suggests that even if the connectivity between VIP^+^ and SST^+^ cells is present at this time point, it seems to be overridden by the overall strong excitation provided by thalamo-cortical inputs onto both IN types. It is interesting to note that the Ca^2+^ transients recorded before P14 upon, especially multi-whisker stimulation, are slower and more prolonged compared to after P14. There are a number of potential mechanisms that could have explained this phenomenon such as action potential speed leading to a slower intracellular Ca^2+^ rise in the cytoplasm or altered buffering and/or extrusion capacities. Nevertheless, we do not detect significant changes over development in the Ca^2+^ transients of SST^+^ INs recorded *in vitro* upon action potential discharge, rather suggesting that reverberating networks could underlie the generation of multiple action potentials spread across time and underlie the slower Ca^2+^ transients. This may be especially the case when multiple whiskers are deflected. In this case, the signal would activate the VPM, which through direct connections activate the VIP^+^ and SST^+^ INs. In addition to this early activation, there would be a later wave of activity onto INs via the engagement of a parallel pathway, consisting of a loop between wS1 and PO^37,38^, leading to the prolonged Ca^2+^ transient component. Nevertheless, our wS1 distance-dependent anatomical and functional connectivity suggests that the cross-barrel circuit is also a contributor to the multi-whisker preference of SST^+^ cells, both before and after P14. We find that excitation coming from lateral cross-barrel domains is stronger onto SST^+^ compared to VIP^+^ cells at both time points, whereas inhibition from within layers 2/3 is stronger onto VIP^+^ cells. This higher inhibition onto the latter cells, could be originating from multiple types of presynaptic GABAergic cells, including other VIP^+^ cells, reelin-expressing interneurons^39^ or even SST^+^ cells, as suggested by Yu et al.^20^

At the same time, we find a striking developmental change in bottom-up thalamic inputs onto SST^+^ INs. Whereas prior to active whisking they receive the majority of inputs from the sensory-related nuclei VPM and PO, after the onset of active whisking their thalamic inputs shift to motor-related nuclei, such as the VM and the VAL. A study by Wall et al., that combined AAV helper virus and monosynaptic rabies virus to trace the inputs onto inhibitory INs in the adult S1, showed that the VPM and PO strongly project to SST^+^ INs, while the motor-related nuclei only showed few presynaptic cells^40^. However, it is important to note that in our experiments we get very few starter cells in the main thalamo-recipient layer (L4) compared to the abovementioned study. Additionally, due the usage of whole brain clearing and the autofluorescence of the thalamic barreloids (VPM), we have been able to very precisely assign cells to different thalamic nuclei, which could have been mis-assigned to more rostral VPM using histological approaches.

Interestingly, another study that reported the in-vivo activity of SST^+^ neurons in adult animals showed that they do not receive strong inputs from the VPM^20,32^. The authors suggested that SST^+^ INs may be recruited rather by local excitatory inputs, which fits well with our cortical connectivity results onto these interneurons. However, we can certainly not exclude that some of the SST^+^ activation comes from different thalamic nuclei such as VM and VAL. These nuclei are in fact known to project to the striatum, L1 of motor-related cortical areas, the cingulate as well as the prefrontal cortex^41–45^. A proportion of VM cells also have axonal collaterals to S1, the target specificity of which was until this study unknown^46,47^. AAV tracings from the Allen Mouse Brain Connectivity Atlas (http://connectivity.brain-map.org/projection) show that axonal projections from the VM exclusively target L1 of the somatosensory cortex. This strongly suggests that the connection we uncover with the rabies tracing is formed between VM and L2/3 SST^+^ cells as they are known to extend their dendrites into L1^48^. Furthermore, the VM is activated during whisking events, as assessed by *in vivo* photometry with GCaMP6^49^, suggesting that it is involved in reporting whisker-related information which it would pass on to the superficial SST^+^ INs.

Notwithstanding the underlying mechanisms, both VIP^+^ and SST^+^ INs show greater responses upon multi-compared to single-whisker stimulation before P14. Together with our anatomical data, this indicates that the functional impact of bottom-up input onto VIP^+^ cells decreases and becomes more specific after P14. We would speculate that this decrease coincides with the emergence of top-down modulation from wM1 to allow these cells to act as coincident detectors between active whisking and touch of individual whiskers. The SST^+^ INs on the other hand would be preferentially activated if multiple whiskers are deflected simultaneously. We would argue that the ability of SST^+^ INs to differentiate between single-and multi-whisker stimuli in the juvenile wS1, suggests that these cells act antagonistically to the activation of VIP^+^ cells, depending on how each whisker is deflected in relation to their neighboring ones. We would speculate that upon complex navigation or object exploration, in some parts of the wS1 SST^+^ activation would take over, while VIP^+^ cells would be more engaged in other parts. Therefore, our results support a model that upon the onset of active whisking, VIP^+^ and SST^+^ INs diversify their functions to be able to register multiple aspects of the environment and hence help facilitate appropriate motor outputs.

## Methods

### Animals

All animal experiments were approved by the Cantonal Veterinary Office Zurich and followed Swiss national regulations. Animal lines used in this study are: VIP-IRES-Cre (Vip^tm1(cre)Zjh^/J)^50^, SST-IRES-Cre (Sst^tm2.1(cre)Zjh^/J)^50^ Ai14 (B6;129S6-Gt(ROSA)26Sor^tm14(CAG-^ tdTomato)Hze/J)51 and HTB (Gt(ROSA)26Sortm1(CAG-neo,-HTB)Fhg)52.

### Animal surgery and preparation for in vivo imaging experiments

We used 6 VIPCre-tdTomato and 6 SSTCre-tdTomato mice (7 males, 5 females) at ages ranging from P8 to P41 for two-photon Ca^2+^ imaging. Mice were sedated with chlorprothixene (0.1g/kg, intraperitoneal (i.p.); Sigma-Aldrich Chemie GmbH, Buchs, Switzerland) and lightly anesthetized with urethane (0.25–0.5g/kg, i.p.). Atropine (0.3 mg/kg; Sigma-Aldrich Chemie GmbH, Buchs, Switzerland) and dexamethasone (2 mg/kg; aniMedica GmbH, Senden-Bösensell, Germany) were administered subcutaneously (s.c.) to reduce secretion of saliva and to prevent edema (s.c. injection 30 min after induction of anesthesia). The body temperature was maintained at 37º C with a heating pad. Hydration levels were checked regularly and maintained by s.c. injections of Ringer-lactate (Fresenius Freeflex; Fresenius Kabi AG, Oberdorf, Switzerland). The depth of anesthesia was evaluated throughout the experiment by testing the pinch reflex on the forepaw. A custom-built head plate was glued to the skull over the left hemisphere with dental cement (Paladur, Heraeus Kulzer GmbH Hanau, Germany; Caulk Grip Cement for electrophysiology) to secure and stabilize the animal.

A small cranial window of 1.5×1.5 mm^2^ was opened above the center of the mapped barrel columns with a sharp razor blade and superfused with Ringer’s solution (in mM: 145 NaCl, 5.4 KCl, 10 HEPES, 1 MgCl_2_, 1.8 CaCl_2_; pH 7.2 adjusted with NaOH). Care was taken not to damage the dura or surface blood vessels in young animals. In animals older than P20, we removed the dura to prevent blockage of the glass pipette tip during insertion into the cortex for two-photon guided Ca^2+^ indicator loading.

### Intrinsic optical imaging

The principal whisker-related barrel column was identified using optical imaging of intrinsic signals. The cortical surface was visualized through the intact bone by surface application of normal Ringer’s solution and a glass coverslip placed on top. The skull surface above the barrel cortex was left intact for animals younger than P12, but thinned in older animals. Reference images of the cortical blood vessel pattern were visualized by a 546-nm LED to enhance contrast. Functional maps of the target barrel column C2 were obtained by shining red light (630 nm LED) on the cortical surface while stimulating the C2 whisker with a galvanometer (10 Hz for 2s at 1140º/s amplitude in rostro-caudal direction^21^). Reflectance images were collected through a 4x objective with a CCD camera (Scientifica SciCam Pro; 14-bit; 2-by-2-pixel binning, 680×512 binned pixels at 31 fps). Functional intrinsic signal images were computed as fractional reflectance changes relative to the pre-stimulus average (average of 10 trials). The intrinsic signal image obtained for the C2 barrel column was then mapped to the blood vessel reference image and used to guide the location of the craniotomy and Ca^2+^ imaging.

### Galvanometer-driven whisker stimulation

Whisker stimulation was performed with a two galvanometer-driven stimulation^21^. One stimulation fiber was attached to the C2 whisker considering variations in resting position angles and relative anterior-posterior shifts. The second stimulator was positioned closer to the whisker pad and a small holder perpendicular to the fiber arm was added for multi-whisker stimulation. For single principal whisker stimulation only the first fiber was moved, whereas for multi-whisker deflection, both the fibers were moved at the same time. The stimulation fibers were fixed and secured with Plasticine on top of a custom-built holder plate and secured and translated with a micro-manipulator. Deflections were applied in the rostro-caudal direction and one single pulse consisting of a phase-shifted 100 Hz cosine with 1140 °/s peak velocity was applied through either one or both galvanometer-driven stimulators.

### In vivo two-photon calcium imaging

Neuronal ensembles in superficial layers of the principal whisker barrel field mapped by intrinsic signal imaging were bolus-loaded with the AM ester form of Oregon Green BAPTA-1 by pressure injection (OGB-1; 1 mM solution in Ca^2+^-free Ringer’s solution; 2-min injection at 150-200 µm depth) as described previously^53^. The craniotomy was then filled with agarose (type III-A, 1% in Ringer’s solution; Sigma) and covered with an immobilized glass plate. Two-photon Ca^2+^ imaging was performed with a Scientifica HyperScope two-photon laser scanning microscope one hour after bolus loading using a Ti:sapphire laser system at 900 nm excitation (Coherent Chameleon; approx. 120 femtosecond laser pulses). Two-channel fluorescence images of 256×128 pixels at 11.25 Hz (HyperScope galvo-mode) were collected with a 16x water-immersion objective lens (Nikon, NA 0.8). 3 to 5 separate spots (i.e. Figure 1C top row) have been imaged per animal. Per imaging spot, both a 300s-long continuous recording of spontaneous activity as well as 10 trials of 20s-long evoked activity recorded for each of single-and multi-whisker stimulation paradigms. Data acquisition was controlled by ScanImage^54^. Duration of Ca^2+^ imaging recordings varied between 3 to 4 hours.

### Analysis of calcium imaging data

Ca^2+^ imaging data were imported and analysed using routines custom-written in MATLAB. First, fluorescence image time-series for a given region were concatenated. The concatenated imaging data was then aligned using a cross-correlation based subpixel registration algorithm^55^ to correct for translational drift (registered on red tdTomato channel and transferred to OGB-1 channel). Average intensity projections of the imaging data were used as reference images to manually annotate regions of interest (ROIs) corresponding to individual neurons. Neurons with somata partly out-of-focus were not included. Ca^2+^ signals were expressed as the mean pixel value of the relative fluorescence change ΔF/F = (*F*-*F*_0_)/*F*_0_ in each given ROI. *F*_0_ was calculated as the bottom 5% of the fluorescence trace. Neuropil patches surrounding each neuron is defined by all pixels not assigned to a neuronal soma or astrocyte of the corresponding neuron ROI annotation^56^(A disk shaped region around the neuron of interest with excluding any intersecting neighboring neuronal ROI). Neuropil correction is performed as F_corrected_ = F_neuron_ – alpha*F_neuropil_. Alpha is estimated for each imaging spot separately using the formula F_blood_vessel_/F_surrounding_neuropil_ ^57^. For each stimulus, the evoked responses of 10 trials were analyzed and the response magnitude expressed as the mean of the evoked ΔF/F integral (%·s; integral of the first 8s response starting at stimulus onset). Pearson’s correlation coefficients of sensory-evoked responses for any two neurons at zero lag were calculated for each single trial evoked calcium traces in a 17.4s window starting from stimulus onset. Spontaneous correlations were calculated by averaging correlations of 1000 randomly segmented 17.4s-long pieces from a 300s long spontaneous recording, to get rid of the trace length dependent correlation fluctuations for the comparison of spontaneous and evoked correlations.

### Decoders

Three different decoders were trained either for the multi-or single-whisker stimulation paradigm, to quantify the decoding capacities of the SSTs and the VIPs at P8-P12 and P21+. The decoders used were random forest, naïve Bayes and Error-correcting output codes (ECOC) that classified a cell type into one out of two groups based on baseline-subtracted average stimulus response trials.^58^ Each classifier was trained 1000 times by randomly setting 70% of the data for training (Matlab function TreeBagger, with parameters *N*_trees_=50, minleaf=5; fitcnb and fitcecoc, respectively). For each trained set, 30% of the remaining data was used to calculate classification accuracy.

### Acute slice electrophysiology

Whole-cell patch-clamp electrophysiological recordings were performed on either SSTCre-tdTomato^+^ or VIPCre-tdTomato^+^ cells in L2/3 of the wS1, in acute slices prepared from P8-12 or P21+ animals.

Animals were anesthetized, decapitated, the brain extracted and transferred to 4°C physiological Ringer’s solution (aCSF), of the following composition (mM): 125 NaCl, 2.5 KCl, 25 NaHCO3, 1.25 NaH2PO4, 1 MgCl2, 2 CaCl2 and 20 glucose. The brain was then glued to a stage and cut into 300 µm-thick coronal slices using a vibratome (VT 1200S, Leica). The slices recovered in room temperature aCSF for 30 min before recording. The slices were then placed in the recording chamber of an upright microscope (Axioscope 2 FS, Zeiss) and superfused with 32°C oxygenated (95% O2 and 5% CO2) aCSF at a rate of 2-3 ml/min. The microscope was equipped with immersion differential interference contrast (DIA) and the following objectives were used to visualize the cells (10x/0.3, Olympus & 40x/0.8, Zeiss). A CMOS camera (optiMOS, QImaging) was attached to the scope to visualize the slice and cells through a computer screen. A white-light source (HAL 100, Zeiss) and a LED based excitation source (Polygon400, Mightex Systems) in combination with a tdTomato filter set (set 43 HE, Zeiss, Excitation 550/25, Emission 605/70) were used to locate the fluorescent INs. Patch pipettes were pulled from borosilicate glass capillaries (1.5 OD x 0.86 ID x 75 L mm, Harvard Apparatus) at a resistance of 4-6 MΩ.

For recordings of intrinsic electrophysiological properties and photostimulation-evoked currents, Clampex was used (v10.7.0.3, Molecular Devices 2016). The recording pipettes were filled with a solution containing the following (mM): 135 potassium D-gluconate, 4 NaCl, 0.3 Na-GTP, 5 Mg-ATP, 12 phosphocreatine-di-tris, 10 HEPES, 0.0001 CaCl2 (pH 7.25, mOsm 290). In all cases 3 mg/ml biocytin (Tocris) was added in the recording solution and no extra pharmacology was added. Access resistance was constantly monitored to ensure the quality and stability of the recording. The recorded data were accepted only if the initial series resistance was less than or equal to 25 MΩ and did not change by more than 20% throughout the recording period. Compensation was made for the pipette and cell capacitance.

We used a similar stimulation protocol as previously reported^59,60^. For each analyzed cell, passive and active membrane properties were recorded in current-clamp mode at −65 mV by applying a series of linearly increasing hyperpolarizing and depolarizing sub-and supra-threshold current steps (500 ms, Δ+20 pA). The analysis of intrinsic properties was done in Clampfit (v10.7.0.3, Molecular Devices 2016). Using this protocol, we analyzed the cells for 5 different parameters: For calculating the membrane resistance, we used the R=V/I on hyperpolarizing pulses that did not lead to an activation of voltage-dependent conductances; Spike threshold was obtained as the inflection point of a rapid change in dV/dt of the first spike evoked at the step protocol. The spike height was from the threshold voltage point to the maximum value of the peak. Spike width was calculated as the width at half of the maximum amplitude; Afterhyperpolarization amplitude was measured from spike threshold to the lowest occurrence point after the repolarization phase of the spike.

All intrinsic electrophysiology data presented in the manuscript are average and all statistical comparisons have been done using a Mann-Whitney U test.

### In vitro Calcium transient and AP correlation

Whole-cell patch-clamping of SSTCre-tdTomato^+^ neurons of L2/3 in wS1 was performed at P8-12 and P21+. Patch pipette was filled with 100 μM OGB-1 and right after breaking in to patched cell, an incubation period of 25 min took place to let the OGB-1 equilibrate within the cell body. A square pulse current was injected giving rise to a set number of APs (#1, 3, 5, 7 & 10), while simultaneously recording the evoked Ca^2+^ transient from the somata, using a Scientifica SliceScope two-photon laser microscope and a laser system at 900 nm excitation. Two-channel fluorescent images of 256×128 pixels at 11.25 Hz (HyperScope galvo-mode) were collected with a 40x water-immersion objective lens. The ΔF/F of the gathered Ca^2+^ responses per a given number of APs were analyzed in terms of the Ca^2+^ transient max amplitude, decay tau and integral. The Averages of these parameters were taken and compared between the two age groups, P8-12 and P21+.

### Glutamate uncaging and data analysis

For photostimulation experiments, whole-cell patch-clamp recordings were performed on randomly selected INs in L2/3 and using a 10x objective lens (Olympus, NA 0.3, 1048 x 1960 µm field of view). For light stimulation, the tissue was digitally divided into 400 subregions making up a grid pattern, spanning all cortical layers. The tissue was submerged in oxygenated aCSF containing caged glutamate (295 µM RuBi-Glutamate, Tocris) and photostimulation was performed at least 3 times at each spot, in a pseudorandom manner. Photostimulation parameters were calibrated beforehand so that the experimental paradigm successfully evoked a short train of ~4 APs in randomly selected putative pre-synaptic partners of the recorded cells. The average time of the evoked APs in putative pre-synaptic neurons were used as a window of analysis for the responses evoked within the patched INs (Figure 3a). This time-window was used to capture majority of mono-synaptic activity and to minimize the presence of poly-synaptic. Although, when performing the glutamate uncaging, the activation of presynaptic partners through photostimulation never resulted in AP elicited currents within the postsynaptic patched cell, suggesting that the contribution of polysynaptic activity would be small. Recordings of synaptic activity were performed in voltage clamp mode at −45 mV (to register both excitatory and inhibitory currents), with a sampling rate of 10 kHz^61–63^.

Custom-written Matlab script was used to analyze the photostimulation evoked current data. Within the data gathered at −45 mV, the excitatory currents were extracted by taking all activity registered below baseline (average of the signal within 500 ms before illumination) within the analysis window, while inhibitory currents were all activity above baseline.

For each recording, the field of view was segmented per layer and the average activity within each layer was normalized to the overall average activity within the whole field of view. Mann-Whitney U test was carried out for all data points within each layer and comparison was made between lamina.

For the lateral activity analysis, the average activity within L2/3 was calculated per given distance from each recorded cells soma. The average was then calculated for all recordings within a given group. Two-tailed t-test was used to look for significance between groups and points with same given distance from soma.

A second set of photostimulation experiments were performed on SSTCre-tdTomato^+^ or VIPCre-tdTomato^+^ cells in L2/3 of the wS1, in acute slices prepared from P8-12 animals. This set of glutamate uncaging experiments was performed in same manner as described above but at −70 mV, to allow to selectively detect only incoming excitatory currents to our recorded cells. When a first round of photostimulation had been carried out, 1 µM TTX (Tetrodotoxin citrate, Tocris) was added to the circulating aCSF and 12 min after, the same glutamate uncaging experiment was performed. Through custom-written Matlab script we then subtracted the values obtained before from after TTX to calculate the extent of the synaptically evoked responses originating from pre-synaptic partners alone, rather than direct stimulation of the recorded cell. When performing this analysis only evoked activity originating from the superficial layers were considered.

### Immunohistochemistry

For the immunohistochemistry experiments, the Ai14 reporter mouse line was combined with either VIPCre or SSTCre. The offspring were sacrificed at either P9-12 (before onset of active whisking) or at >P21 (after the onset of active whisking). In short, the animals were anesthetized before being transcardially perfused with ice-cold 1x PBS followed by 4% PFA. The brains were then dissected and post-fixed in ice-cold 4% PFA for 1h before being placed in a 30% Sucrose solution at 4°C for >24h. Following this cryo-protection step the brains were embedded in OCT using a peel-away mold and then stored at −80°C. Coronal 20μm-think brain sections that contained the barrel cortex were cut and collected on-slide (wS1) using a cryostat (Leica, CM3050 S). The slides were then stored at −80°C until further processing. For labeling thalamic input onto the tdTomato expressing cells an antibody staining was performed. For blocking, 0.75uL of PBS with 0.1% TritonX and 1.5% normal donkey serum was applied on the slide and left for 1h at RT. Both primary and secondary antibodies were diluted in the same blocking solution. The primary, rabbit αVGlut2 (1:500, SySy 135 402), was left on the slide overnight (>17h) at 4°C while the secondary, donkey αrabbit488 (1:1000, Thermo Fisher A21206), was left for 2h at RT. The slides were then coverslipped using VectaShield Gold and stored at 4°C for imaging.

### Confocal imaging and image processing of immunohistochemical samples

The slides were imaged using a Confocal Microscope (Olympus FV1000). All images were taken from L2/3 of the barrel cortex and of each of the processed brain 8-10 stacks were generated. To image the VGlut2 puncta a 60x oil-immersion objective was used and stacks were taken at 0.47μm distance.

A custom MATLAB script was used to analyze the VGlut2-tdTomato colocalization. The algorithm worked as follow: Image de-noising was performed using Wiener filtering followed by 2-D bilateral filtering. Wiener filter performs 2-D adaptive noise-removal resulting in a low pass filtered version of an intensity image that has been corrupted by stationary additive noise^64^ that is estimated based on statistics at a local neighborhood of each pixel.

The bilateral filter is an edge-preserving nonlinear filter that smoothens a signal while preserving strong edges^65^. After de-noising, Z-stacks were binarized using multilevel Otsu thresholding^66^ for each xy-, xz- and yz-slices separately, resulting in 3 binary 3D-stacks combined using logical AND operator (This very stringent binarisation operation makes sure the detected signal is noise free). Finally 3D binarized stacks for channel 1 (VGlut2) and 2 (VIP-or SST-tdTomato) were combined with another AND operator and 3D connected components analysis^67^ was used to count number of co-localizations. Number of co-localizations was normalized by total tdTomato signal for each image.

### Viral injections

For these experiments the HTB reporter line was crossed with either VIP-Cre or SST-Cre. The pups were injected with ASLV-A envelope glycoprotein (EnvA)-pseudotyped, glycoprotein-deleted rabies virus SADG-mCherry(EnvA)^26^ at either P4/P5 or P15. All pups were injected with 100nl of the virus, 120-170um deep into the barrel field of the primary somatosensory cortex. The injections were done using a glass micropipette attached to a Nanolitre 2010 pressure injection apparatus (World Precision Instruments). After the surgeries, the animals were returned to their home-cage for 7 days to allow for adequate viral expression.

For the control rabies virus experiments VIPCre pups aged P15 were injected with 100nl of a 1:1 mix of pAAV-hSyn-FLEX-TVA-P2A-EGFP-2A-oG (gift from John Naughton, Addgene plasmid # 85225) and rabies virus (see above). The injections were done at the same depth and using the same equipment as described above. The animals were sacrificed 7days after the injection and all the brains were cleared as described below.

### Tissue processing after viral injections

Seven days after the rabies virus injection the animals were transcardially perfused using 4% PFA. Following that the brain were either processed for immunohistochemistry as described above or for tissue clearing. For immunohistochemistry the whole brain was cut at 20um using a cryostat (Leica, CM3050 S) and collected on slide. Every third of the collected sections was stained with GFP (abcam, ab13970, 1:1000) to enhance the HTB signal.

The method used for hydrogel-based tissue clearing is described in detail elsewhere^68–70^. In short, the brains were post-fixed for 48 hours in a Hydrogel solution (1% PFA, 4% Acrylamide, 0.05% Bis)^68,69^ before the Hydrogel polymerization was induced at 37°C. Following the polymerisation the brains were immersed in 40mL of 8% SDS and kept shaking at room temperature until the tissue was cleared sufficiently (10-40 days depending on the age of the animals). Finally, after 2-4 washes in PBS, the brains were put into a self-made refractive index matching solution (RIMS)^71^ for the last clearing step. They were left to equilibrate in 5mL of RIMS for at least 4 days at RT before being imaged.

### Imaging of rabies injected brains

A Slidescanner (Zeiss, AxioScan Z1) was used to image the stained sections. Mosaic images of the injected hemisphere were taken using a 20x objective. The obtained pictures were processed for analysis using the ZEN Software and Fiji.

After clearing, brains were attached to a small weight and loaded into a quartz cuvette, then submerged in RIMS and imaged using a home-built mesoscale selective plane illumination microscope mesoSPIM^27^. The microscope consists of a dual-sided excitation path using a fiber-coupled multiline laser combiner (405, 488, 515, 561, 594, 647 nm, Omicron SOLE-6) and a detection path comprising an Olympus MVX-10 zoom macroscope with a 1x objective (Olympus MVPLAPO 1x), a filter wheel (Ludl 96A350), and a scientific CMOS (sCMOS) camera (Hamamatsu Orca Flash 4.0 V3). For imaging mCherry and eGFP a 594 nm excitation with a 594 long-pass filter (594 LP Edge Basic, AHF) and 488 nm & 520/35 (BrightLine HC, AHF) were used respectively. The excitation paths also contain galvo scanners (GCM-2280-1500, Citizen Chiba) for light-sheet generation and reduction of streaking artifacts due to absorption of the light-sheet. In addition, the beam waist is scanning using electrically tunable lenses (ETL, Optotune EL-16-40-5D-TC-L) synchronized with the rolling shutter of the sCMOS camera. This axially scanned light-sheet mode (ASLM) leads to a uniform axial resolution across the field-of-view of 5-10 µm (depending on zoom & wavelength). Field of views ranged from 10.79 mm at 1.25x magnification (Pixel size: 5.27 µm) for overview datasets to 3.27 mm at 4x (Pixel size: 1.6 µm). Further technical details of the mesoSPIM are described elsewhere (www.mesospim.org). The images generated with the mesoSPIM were further processed using the Fiji and the Imaris software.

### Analysis of cleared tissue imaging data

Close-up stacks taken from the barrel cortex of rabies injected and cleared SSTCre-HTB and VIPCre-HTB brains were initially processed through ImageJ and z-stacked samples of superficial and deeper layers were saved after intensity thresholding. In each brain, barrels were segmented from layer 4 by drawing the vectors on barrels through scalar vector graphics’ software (BoxySVG). Afterwards, automated neuron (starter- and presynaptic cells) detection was performed on maximum intensity projections of L2/3 green (HTB) and red (Pre-synaptic partners) through a Deep Neural Network-based method, DeNeRD ^28^. Initially, selected brain sections of high-resolution are segmented into smaller images by equally dividing the brain section into 100×100 smaller sections, whereas each smaller image contains ~20 neurons. After annotating the neurons in a subset of these images (total=160), 2/3rd of them are randomly drawn for training and 1/3rd for testing purpose. A Simple Graphical User Interface (SiGUI) software, developed in MATLAB, is used to generate the ground-truth labels. Ground truth labels were generated by human experts who annotated the bounding boxes on top of the neurons. After generating the annotation, these brain images were fed to the DeNeRD for training the deep neural network (DNN). Four steps training procedure is applied that includes: (i) Training of the Region Proposal Network (RPN), (ii) Use the RPN from (i) to train a fast RCNN, (iii) Retraining RPN by sharing weights with fast RCNN of (ii). Finally, (iv) Retraining Fast RCNN using updated RPN. Epoch size of 500 is used with initial learning rate of 1×10-5 of 1×10-6 in stages (i-ii) and (iii-iv) respectively. Training is performed by using NVIDIA Quadro P4000 GPU. The network is trained by minimizing the multi-task loss which corresponds to the each labeled Region of Interest, ROI (i.e. neuronal body) through stochastic gradient descent algorithm. The average precision (F1 score) of 0.9 was achieved on the testing dataset. After the DNN is trained, each brain section from the barrel region is serially passed through the DNN and neuron segmentation is performed.

Then, the 2D Euclidean distance between each starter cell and each of the pre-synaptic cells present within a 800μm distance was measured using a custom MATLAB code. Plotted was the cumulative distribution of the distance for every starter cell as well as the average cumulative distribution of the percentage (probability) of pre-synaptic partners at a given Euclidian distance from the center of starter cells.

Additional analysis was performed by randomizing the selection of starter and presynaptic cells in the barrel cortex images of each cleared brain. We started by taking the original number of starter cells and randomly selecting the number of presynaptic cells that matches the observed ratio between the two cell groups for each brain. Ten iterations were performed and the Euclidean distance between each starter and presynaptic cell was measured. Subsequently, the number of starter cells was decreased by one and the above process was repeated until only one starter cell was left. For each iteration, the cumulative distance distribution was calculated and plotted.

### Statistical analysis

Data are represented as mean ± s.e.m. unless stated otherwise. Statistical comparisons have been done using a paired Wilcoxon-signed rank test for paired data and two-tailed Mann-Whitney *U*-test for non-paired data. A Kolmogorov-Smirnov test was applied to statistically compare the cumulative distribution of the distance between starter cells and pre-synaptic partners. Two-tailed t-test was used to analyze the distance analysis of the glutamate uncaging evoked responses in L2/3. Significance threshold was set to *p*<0.05; in the figures, different degrees of evidence against the null hypothesis are indicated by asterisks (*p*<0.05: *; *p*<0.01: **; *p*<0.001: ***).

## Data and Code availability

Data and Custom written codes are available upon reasonable request.

## Acknowledgement

We thank O. Hanley for the production of the pseudotyped rabies virus used in this study, A. Engmann and the Salk Institute for providing the cell lines for this production, P. Bethge for his help with the 2P setup, H. Kasper and M. Wieckhorst for technical assistance and O. Weinmann for his advice on immunostainings. We would also like to thank L. Egolf for her assistance with the VIPCre, SSTCre and Ai14 mouse lines. Slidescanner imaging and data analysis was performed with equipment maintained by the Center for Microscopy and Image Analysis (ZMB), University of Zurich. We thank C. Aemisegger at the ZMB for her assistance with the Slidescanner. This work was supported by grants from the European Research Council (ERC, 679175, T.K and 670757, F.H), the Swiss National Science Foundation (SNSF, 31003A_170037, T.K), Fond zur Förderung des Akademischen Nachwuchs of the UZH Alumni (T.K) and the Swiss Foundation for Excellence in Biomedical Research (R.K and T.K)

## Author Contributions

R.K. performed the anatomical mapping, tissue clearing, immunohistochemistry and wrote the manuscript; R.V. performed functional electrophysiology experiments, *in vitro* Ca^2+^ imaging, contributed to the analysis of *in vitro* electrophysiological data and wrote the manuscript; A.v.d.B. performed in vivo Ca^2+^ imaging experiments and initial virus injections; A.O.A. performed the analysis of the *in vivo* Ca^2+^ imaging data, puncta co-localization in histology, analysis of the *in vitro* electrophysiology data and decoder analysis; A.I. did the analysis of the rabies-traced cells and the their distribution in the cortex. F.F.V. built, provided access to and guidance for the light-sheet microscope, F.H. provided access to the Light-Sheet Microscope and a two-photon microscope. D.K. and A.A. provided initial guidance on tissue clearing. T.K. conceptualized the study, designed the experiments and wrote the manuscript.

## Additional Information

Supplementary Figures S1-S7

Supplementary Movies S1-S8

The authors declare no competing interests

## Notes

### Competing Interest Statement

The authors have declared no competing interest.

