## Supplementary Figures for "Developmental Divergence of Sensory Stimulus Representation in Cortical Interneurons"

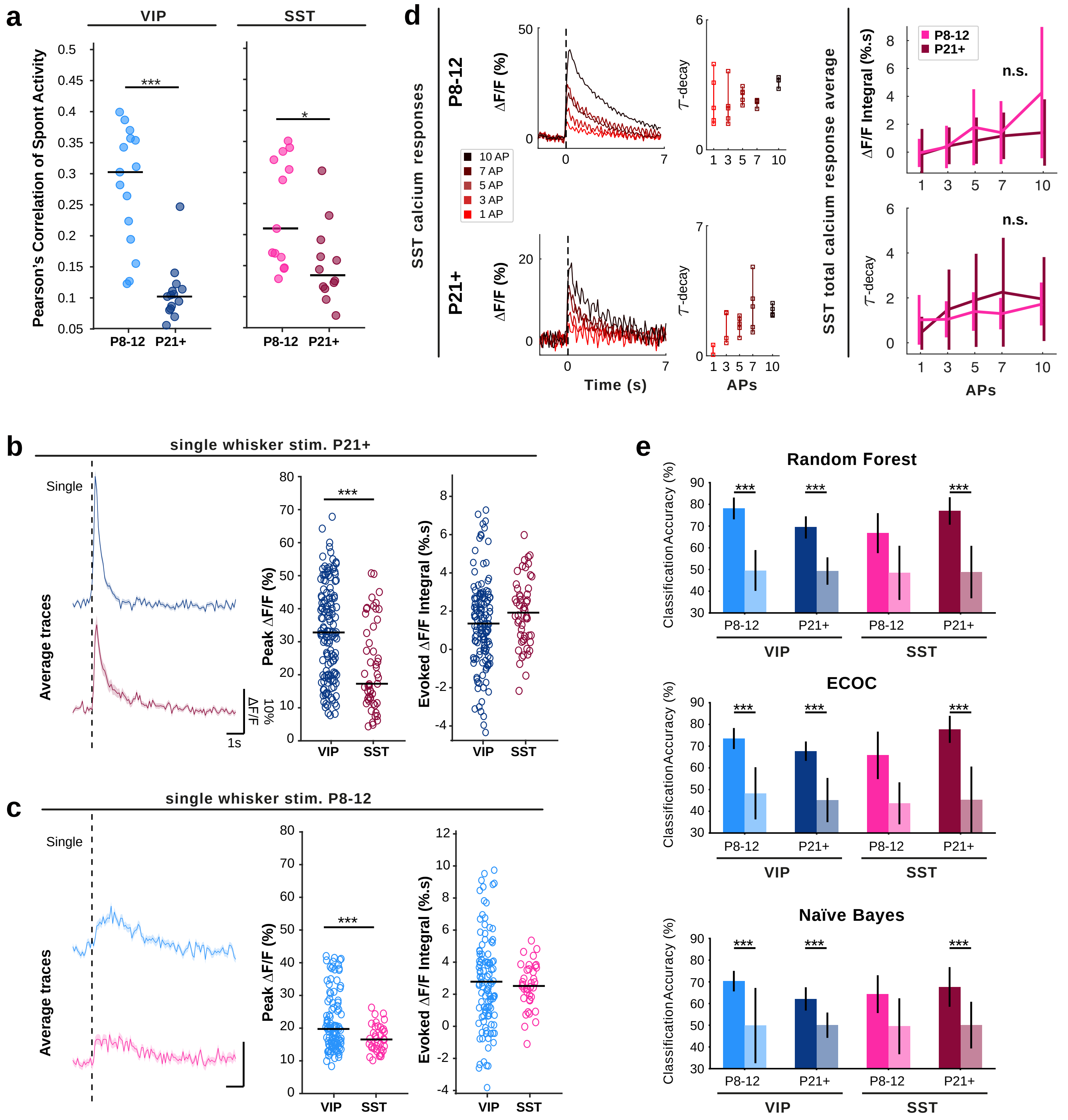


**Supplementary Figure 1: Spontaneous correlation and single whisker stimulation**

**(a)** Pearson's correlation of spontaneous activity between interneurons and the surrounding non-tdTomato expressing cells. **(b,c)** $Ca^{2+}$ response of VIP and SST interneurons to single whisker stimulation at P21+ and at P8-12. Left: Average ΔF/F trace of single whisker evoked activity with SEM shown. Middle: Peak ΔF/F of single whisker evoked activity. Right: Average of the evoked ΔF/F integral. **(d)** Selected examples of $Ca^{2+}$ transients elicited through a set number of evoked APs (1, 3, 5, 7 & 10), within SST interneurons across development. Top: (Left) Averages of ΔF/F calculated from the AP evoked $Ca^{2+}$ transients, in correlation to number of evoked APs. (Right) Tau decay calculated for each ΔF/F average and plotted against number of APs. Bottom: Overall average of ΔF/F (Left) and Tau decay (Right), statistically compared per number of evoked APs across development. (N=3 animals per group, SST P8-12: 4 cells, SST P21+: 5 cells). **(e)** Three different decoders trained (random forest, naïve Bayes & ECOC) for each group (single- and multi-whisker), to see the classification accuracy between stimulation paradigm. Decoding capacities of each interneuron type across development statistically compared against shuffled data. (N=3 animals per group, VIP P8-12: 109 cells, SST P8-12: 38 cells, VIP P21+: 138 cells, SST P21+: 51 cells). Statistics: Mann–Whitney U-test (*p<0.05, **p<0.01, ***p<0.001).

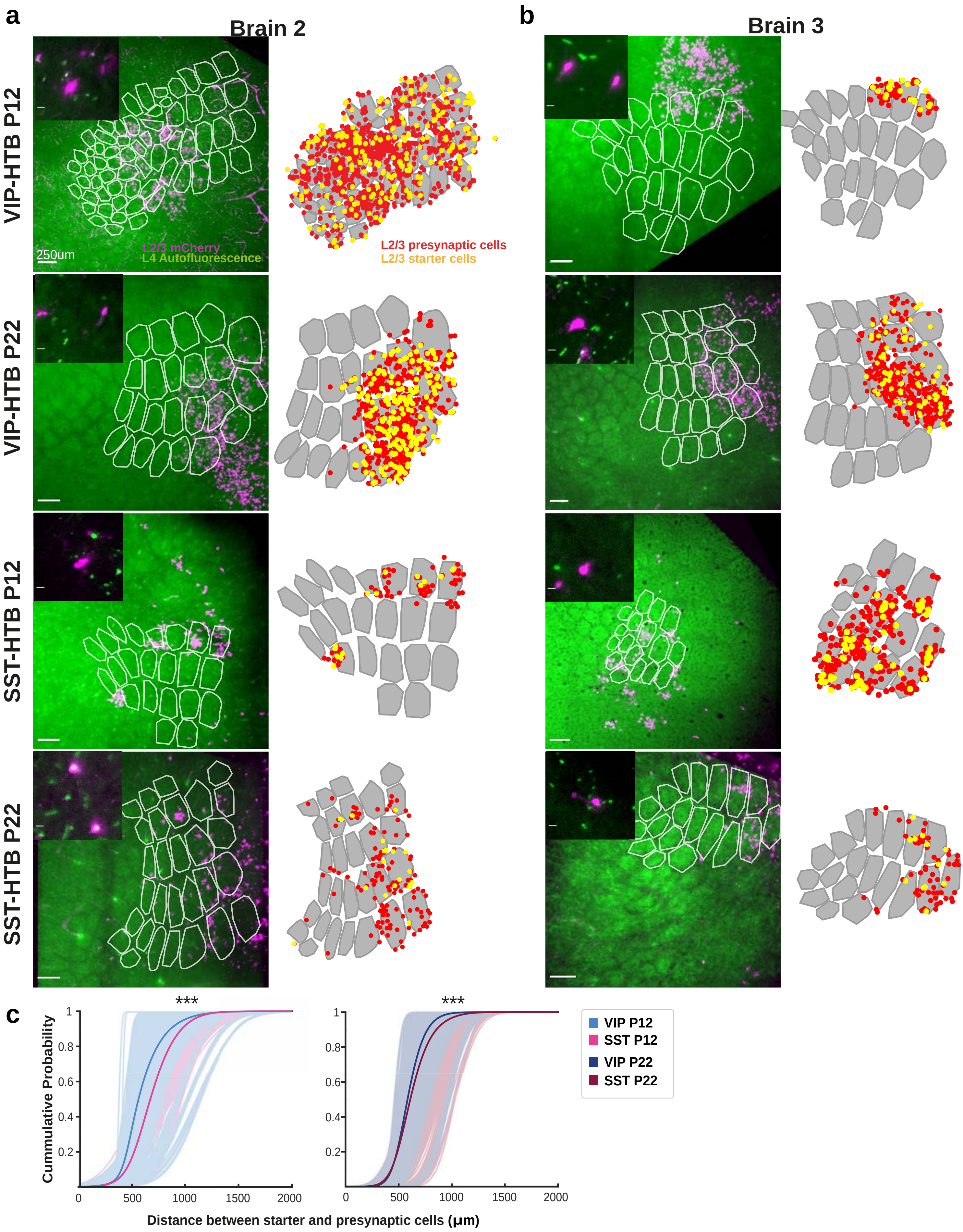


**Supplementary Figure 2: Segmented starter and presynaptic cells for all brains used in the analysis**

**(a&b)** All injection sites included in the analysis. Brain 1 of each genotype and age group is displayed as example in Fig2. Right: Overlay of maximum intensity projection of L2/3 mCherry signal and median intesity projection of L4 Autofluorescence. Inset indicates location of barrel field in the whole brain. Left: Tranformation applied before distance analysis. Segmented barrels overlayed with L2/3 starter (yellow) and presynaptic (red) cells. Insets show close-up of rabies positive neurons.
**(c)** Cummulative Euclidian distance distribution between randomly selected starter and presynaptic cells. Statistics: Kolmogorov-Smirnov Test (***p<0.001).

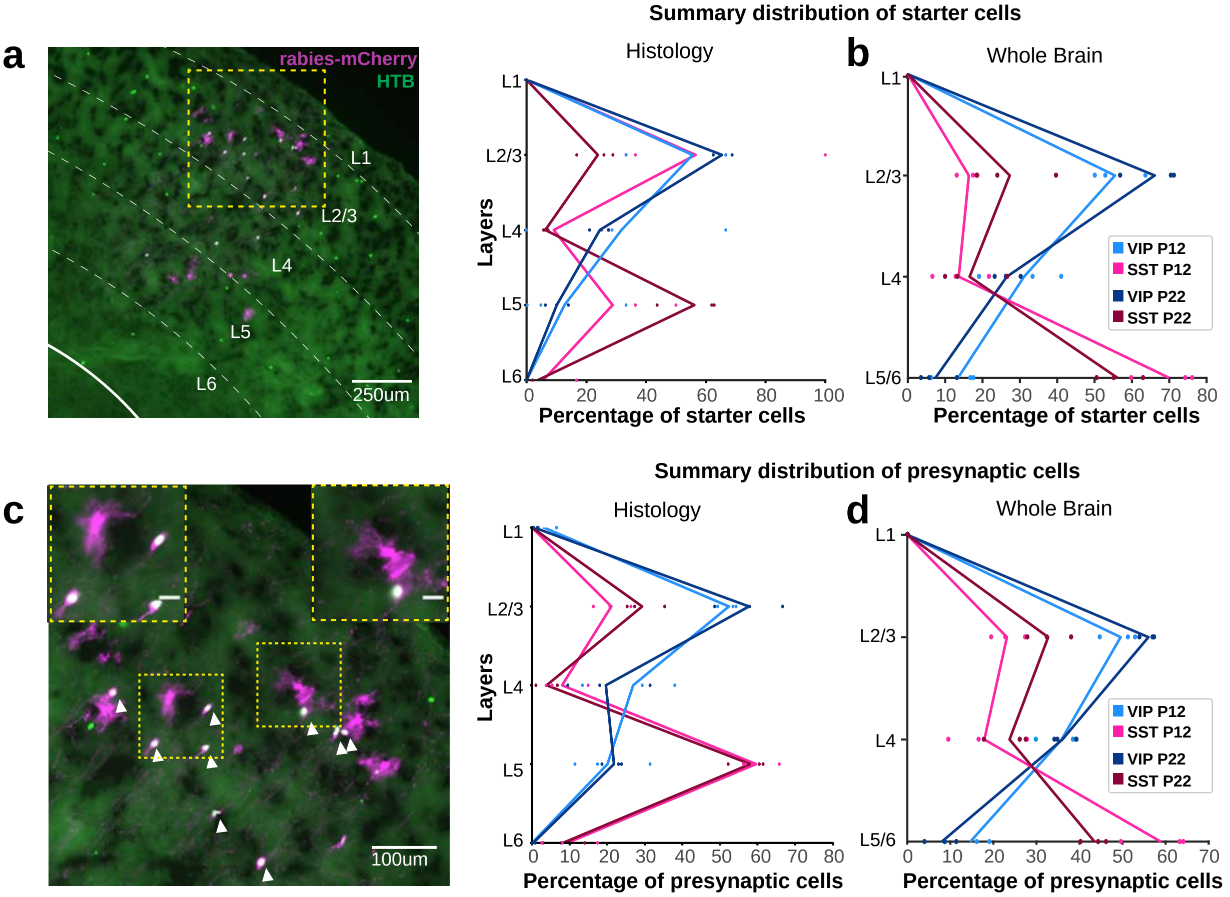


**Supplementary Figure 3: Laminar distribution of starter and presynaptic cells in rabies-injected brains**

**(a)** Left: representative picture of laminar distribution of rabies positive cells in wS1. Right: Distribution of starter cells around the injection site as quantified using a histological approach. **(b)** Distribution of starter cells around the injection site as quantified in the whole brain using a deep neural network. **(c)** Left: representative picture of starter (arrowheads) and presynaptic cells in L2/3 in the wS1 of a VIPCre-HTB mouse at P22. Insets show close-up of rabies positive neurons. Right: Distribution of presynaptic cells around the injection site as quantified using a histological approach.**(d)** Distribution of presynaptic cells around the injection site as quantified in the whole brain using a deep neural network.

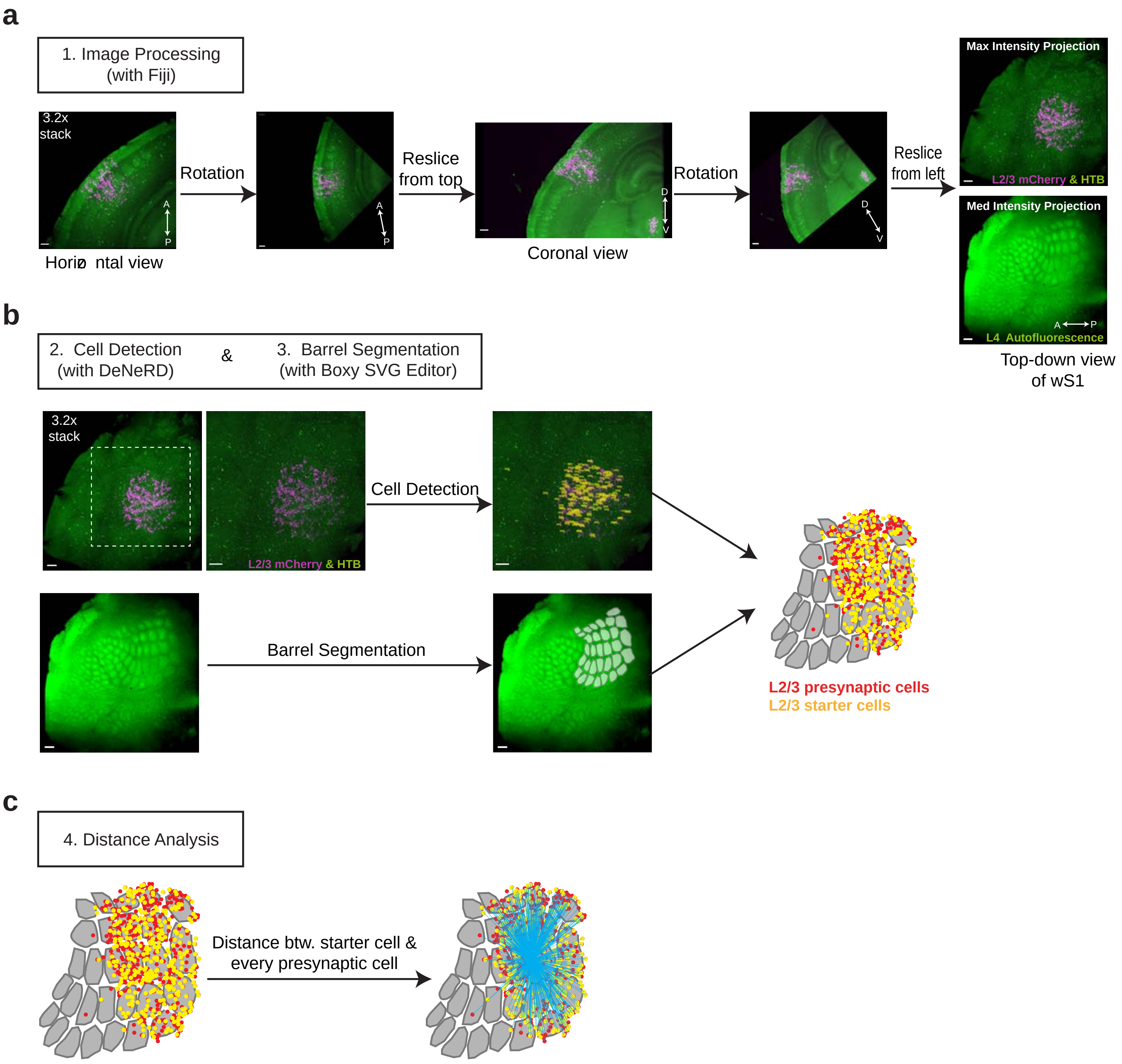


**Supplementary Figure 4: Image Analysis Method**

**(a)** Schematic representation of image processing step. Images are maximum intensity projections, but processing is applied to the entire stack **(b)** Schematic represenation of cell detection and barrel segmentation. The images are passed through a deep neural network (termed DeNeRD) which is trained to detect cells. The resulting output detects starter and presynaptic cells and their locations. Barrels are segmented manually using an SVG editor. Detected dells are overlaid with the segmented barrels to better visualize their location. Cells located outside of barrel field are not included in the analysis. **(c)** Schematic representation of the analysis on the detected cells. Distance of every starter to every presynaptic cell in a 800um radius around it is measured.Blue lines indicate the distance of one starter cell to every presynaptic cell.

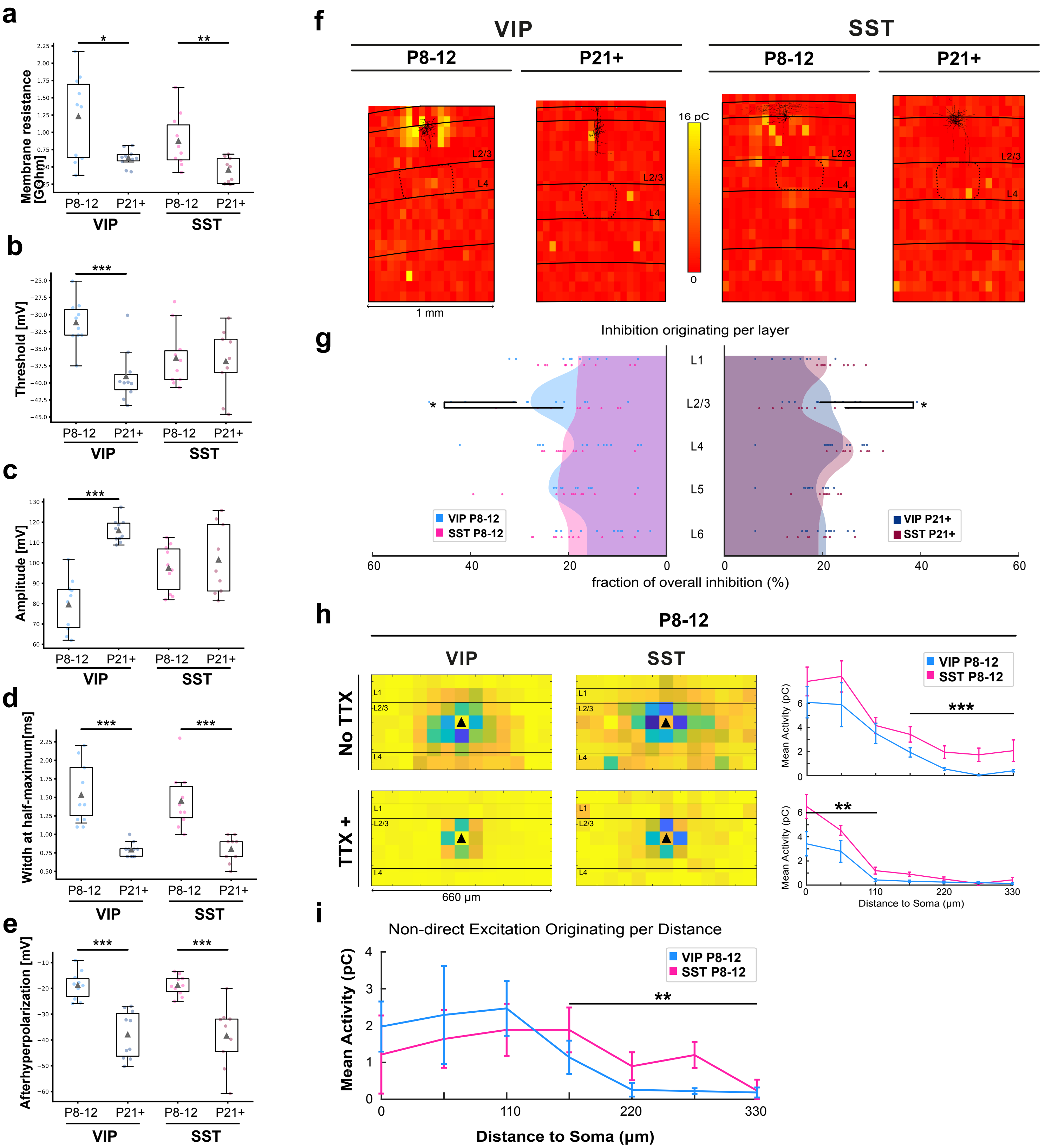


**Supplementary Figure 5: Intrinsic electrophysiological properties of developing intereneurons and laminar origin of incoming inhibition**

**(a-e)** Membrane resistance, threshold for action potential generation, amplitude, width at half-maximum of action potential and afterhyperpolarization from the four groups represented as box plots with each cell shown as a dot and SD bars included. **(f)** Heatplot representations of normalized evoked inhibitory current integral (in pC), recorded over development (P8-12 and p21+) from VIP (left) and SST (right) interneurons while performing glutamate uncaging in a grid pattern. **(g)** Plot of inhibitory input onto each VIP and SST cells (individual dots), averaged per lamina and normalized to average overall inhibition within the field of view. The grand average of all cells per group is depicted as a continuous filled wave and compared within age groups. **(h)** Heatplot representations of overall evoked excitatory current integral (in pC), recorded at P8-12 from VIP (left) and SST (middle) interneurons, recorded without (top heatplots) and with TTX (bottom heatplots). Mean evoked excitation originating from within L1-3 plotted as a function of lateral distance from either side of the recorded cells somata (right), either without (right, top) or with TTX (right, bottom). **(i)** Overall activity recorded in presence of TTX subtracted from activity gathered without. Remaining non-direct activity plotted as a function of lateral distance from either side of the recorded cells somata. (VIP P8-12: 10 cells, SST P8-12: 10 cells, VIP P21+: 10 cells SST P21+: 9 cells). Statistics: Mann-Whitney U test, (*p<0.05, **p<0.01, ***p<0.001).

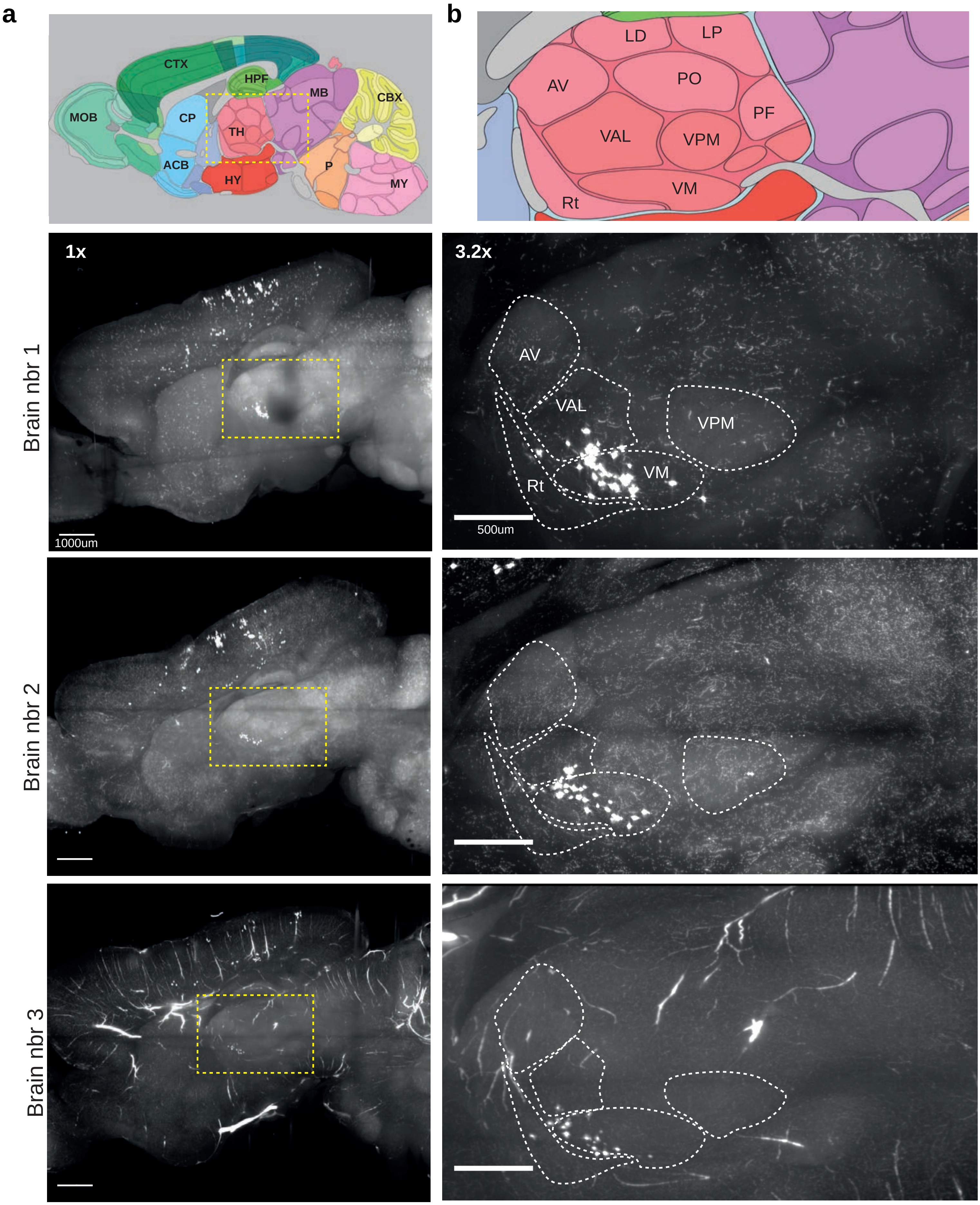


**Supplementary Figure 6: Presynaptic cells in the VM**

**(a)** Overview of all SSTCre-HTB brains injected with rabies at P15 and sacrificed at P22. Top panel: reference atlas from http://atlas.brain-map.org/. Images are sagittal maximum intensity projections. Note: brain nbr 1 is also displayed in Fig4. Also note: much of the fluorescence signal in brain nbr3 comes from blood vessels. This is an artifact of a suboptimal perfusion. Blood vessels can be clearly differentiated from cells based on their shape. **(b)** Zoom-in of thalamus showing the different thalamic nuclei that contain presynaptic cells. Rt: Reticular nucleus, AV: Antroventral nucleus, VAL: Ventral anterior-lateral comples, VM: Ventral medial nucleus, LD:Lateral dorsal nucleus, PO: Posterior complex, VPM: Ventral posteriormedial nucleus, LP: Lateral posterior nucleus, PF: Parafasicular nucleus.

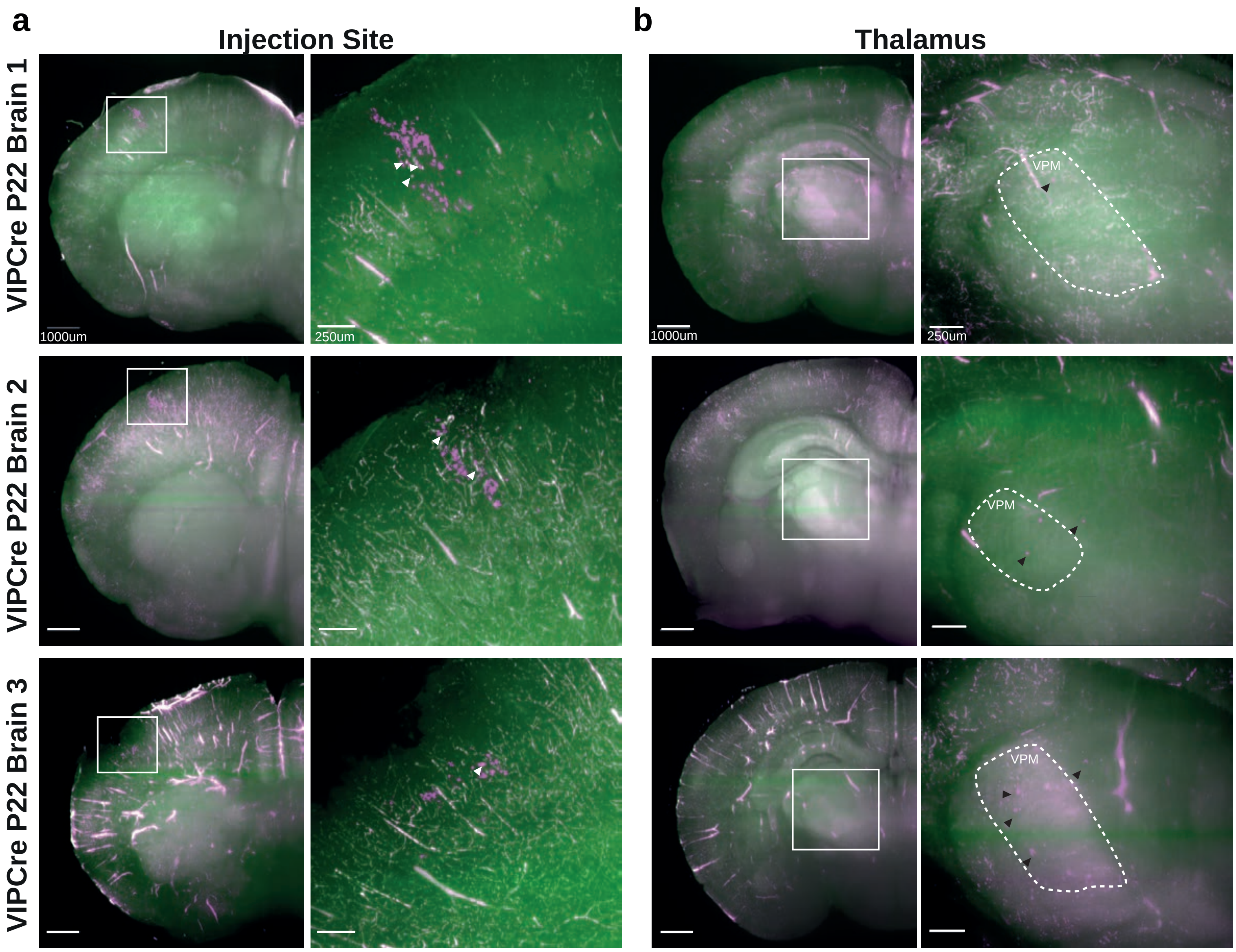


**Supplementary Figure 7: VIP+ cells show presynaptic cells in the thalamus at P22 when the rabies virus is combined with a helper virus**

**(a)** Left: Overview of injection site in 3 cleared brains. Right: zoom-in of the injection site with starter cells (white arrowheads). Note: not all starter cells are depicted in this maximum intensity projection. **(b)** Left: overview of the thalamus. Right: zoom-in of the thalamus, showing the Ventral posteriormedial nucleus (VPM) with the rabies infected presynaptic cells (black arrowheads). Images are maximum intensity projections.
